# Tenascin-C from the tissue microenvironment promotes muscle stem cell self-renewal through Annexin A2

**DOI:** 10.1101/2024.10.29.620732

**Authors:** Mafalda Loreti, Alessandra Cecchini, Collin D. Kaufman, Cedomir Stamenkovic, Alma Renero, Chiara Nicoletti, Anais Kervadec, Gabriele Guarnaccia, Daphne Mayer, Alexandre Colas, Pier Lorenzo Puri, Alessandra Sacco

## Abstract

Skeletal muscle tissue self-repair occurs through the finely timed activation of resident muscle stem cells (MuSC). Following perturbation, MuSC exit quiescence, undergo myogenic commitment, and differentiate to regenerate the injured muscle. This process is coordinated by signals present in the tissue microenvironment, however the precise mechanisms by which the microenvironment regulates MuSC activation are still poorly understood. Here, we identified Tenascin-C (TnC), an extracellular matrix (ECM) glycoprotein, as a key player in promoting of MuSC self-renewal and function. We show that fibro-adipogenic progenitors (FAPs) are the primary cellular source of TnC during muscle repair, and that MuSC sense TnC signaling through cell the surface receptor Annexin A2. We provide in vivo evidence that TnC is required for efficient muscle repair, as mice lacking TnC exhibit a regeneration phenotype of premature aging. We propose that the decline of TnC in physiological aging contributes to inefficient muscle regeneration in aged muscle. Taken together, our results highlight the pivotal role of TnC signaling during muscle repair in healthy and aging skeletal muscle.

## INTRODUCTION

Muscle stem cells (MuSC) are a muscle resident stem cell population responsible for developmental and postnatal muscle growth as well as adult tissue repair ^1^. In healthy adult conditions, MuSC reside in a quiescent state in their sublaminar niche, located between the muscle fiber membrane and the basal lamina. In response to damage, MuSC become activated and proliferate to then either replenishing the stem cell pool (self-renewal) or differentiating and repairing the injured tissue (reviewed in ^2^). Each stage of the MuSC differentiation process, called myogenesis, occurs in a timely and spatially defined manner to ensure proper muscle regeneration and a full restoration of its function.

The tissue microenvironment is a crucial regulator of muscle maintenance and repair ^3^. The MuSC milieu contains a heterogenous population of cells (including fibro-adipogenic progenitors (FAPs), endothelial cells, muscle-resident immune cells), extracellular matrix (ECM) proteins (*e.g*., collagen, laminin, tenascin-C (TnC)) and growth factors ^3^. Among the tissue resident cells, FAPs play a pivotal role in guiding MuSC-mediated repair. It has been previously shown that FAPs assist the regeneration process by promoting MuSC differentiation ^4,5^. Muscle-specific ablation of FAPs results in a muscle that is unable to fully regenerate ^6^. FAPs secret signaling molecules and deposit ECM components to promote muscle maintenance and repair and provide support to the MuSC niche ^7-12^. Although recent studies have provided evidence of a major role of the crosstalk between FAPs and MuSC in skeletal muscle, we still have a limited understanding of the local cues that mediate these interactions.

The extracellular matrix (ECM) provides not only structural support for the physical localization of quiescent MuSC, but ECM proteins also mediate signaling through direct interaction with MuSC cell surface receptors ^13^. In chronic conditions (*e.g*., aging, myopathies), changes in ECM composition can impair MuSC self-renewal and regeneration, impacting tissue maintenance ^14^. Our previous findings, together with work carried out by other laboratories, show that the ECM glycoprotein TnC is expressed by MuSC during both embryonic development and adult tissue repair ^15-18^. In skeletal muscle, TnC is widely expressed during embryonic development, while in adulthood its expression is only transitional during regeneration or it is restricted to defined regions, such as tendons and ligaments ^18,19^. We have previously demonstrated that transplantation of TnC-downregulated fetal MuSC in injured muscle reduced their contribution to tissue repair, while this effect was not observed in the adult control (TnC-downregulated adult MuSC) ^18^. TnC can also be released by necroptotic myofibers upon injury and is important for coordinating muscle regeneration through increased MuSC proliferation ^20^. TnC is also abundantly expressed and secreted by glial cells near NMJs upon muscle denervation ^21,22^. These data confirm that TnC is produced by multiple cellular sources within the damaged adult skeletal muscle microenvironment to facilitates muscle repair ^18,23^. It has been shown that genetic deletion of TnC induces behavioral abnormalities (hyperlocomotion, coordination defects), delayed olfactory detection during development, impaired angiogenesis, reduced hematopoietic stem cell function, defects in wound healing and muscle atrophy upon mechanical stress ^24-32,33^.

TnC accomplishes its different roles via signaling through multiple binding partners in a context-dependent manner. These include cell surface receptors (*e.g.,* Annexin A2, Syndecan-4, EGFR, TLR4, integrins ⍺5β1/3/6, ⍺2/7/8/9β1), ECM components (*e.g.,* collagen, fibronectin), and soluble factors (*e.g.,* FGF, TGFβ, VEGF, BMP) ^34,35^. Previous studies have demonstrated that TnC shows high-affinity to bind the Ca^2^-dependent phospholipid-binding protein Annexin A2 across many cell types (*e.g.*, cancer, endothelial cells, fibroblasts) and in different species (*e.g*., mouse, human) ^36,37^. TnC:Annexin A2 interaction has been shown to enhance cell migration in diverse tissues ^37,38^. Annexin A2 is also involved in membrane repair of damaged muscle fibers, which makes Annexin A2 a potential target for ameliorative treatments in patients affected by inflammatory myopathies ^39-41^. Although much is known about the effect of the interactions between TnC and cell surface receptors, the exact molecular mechanisms by which TnC signals to myogenic cells in the context of muscle maintenance and repair are still poorly understood.

There is extensive evidence of defective muscle regeneration occurring during aging, due to a disrupted stem cell niche and premature activation of MuSC ^42^. Changes in the ECM composition with age, such as increased stiffness and abnormal signaling from the microenvironment, cause the dysregulation of MuSC quiescence and activation. This dysregulation has negative effects on MuSC proliferation and differentiation, which leads to an overall diminished the regenerative capacity of the tissue ^14,43,44^. These age-related ECM changes also correlate with altered function of other resident cell populations, such as FAPs. During aging, FAPs exhibit reduced proliferation and secretion of signaling molecules, in favor of fibrogenic differentiation ^45,46^. While several signaling networks regulating skeletal muscle regeneration have been previously reported, the cellular and molecular mechanisms underlying TnC signaling in adult or aged skeletal muscle repair have not been investigated.

Here we show that muscles lacking TnC exhibited impaired MuSC self-renewal, had reduced numbers of MuSC, and favored an increased number of committed myogenic progenitors. TnC-ablated MuSC also demonstrated impaired migration, that is rescued by soluble TnC treatment. We further provided evidence that FAPs are the predominant source of TnC during regeneration and that MuSC sense TnC through the cell surface receptor Annexin A2. Finally, we observed that young TnC-KO mice recapitulate the impaired regeneration phenotype of old wild-type mice following injury. Our results suggest TnC and its receptors on MuSC are potential candidates for pharmaceutical interventions to enhance muscle repair in aged tissue by restoring the signaling to a youthful state.

## RESULTS

### Tenascin-C (TnC) regulates the quiescence and commitment of postnatal MuSC

We utilized Tenascin-C knockout (TnC-KO) mice ^47^ to evaluate the role of TnC in skeletal muscle. Constitutive genetic deletion of TnC did not impair skeletal muscle formation during embryogenesis and postnatal maturation. Tibialis anterior (TA) muscles from TnC-KO mice did not have significantly different weight (**Figure 1A**) or myofiber cross-sectional area (**Figure 1B**) compared to wild type (WT) controls in young adulthood (3-6 months of age). However, we did observe a significant reduction in the number of MuSC in the TA muscles of adult TnC-KO compared to age-matched WT controls (**Figure 1C**). We next performed histological analyses of TA muscles at E16.5, P14, and P30 to investigate whether the decline in MuSC numbers in TnC-KO mice arises prenatally, postnatally, or in young adults. We observed that MuSC numbers are unaffected by TnC deletion at P14, while there was a ∼50% reduction at P30, which was consistent with what we observed in adulthood (**Figure S1A**, **Figure 1C**). While there was an initial decrease in TA myofiber CSA at P14, the difference was no longer detectable by P30 (**Figure S1B**), further recapitulating what we detected in adulthood (**Figure 1C**). Previous work has shown that as tissues reach homeostasis during postnatal muscle growth, a greater proportion of MuSC progressively enter quiescence ^18,48-51^. Thus, we investigated whether the decline in MuSC numbers in TnC-KO mice was due to an impairment in their ability to regulate quiescence at this stage. While TnC-KO MuSC proliferation is not affected at any stage (**Figure S1C-F**), we did observe an increase in the number of MyoD^+^ cells in P14 TnC-KO muscles compared to WT controls (**Figure 1D**), indicating a bias towards myogenic commitment. We further observed that P14 TnC-KO muscles exhibited an increase in the percentage of Pax7^+^ MuSC in the interstitial space compared to WT controls, suggesting that TnC plays a role in the proper localization of MuSC (**Figure 1E**). Recent studies have provided evidence that the quiescent state of MuSC is characterized by the presence of cellular projections, which are lost during activation ^52,53^. We investigated whether TnC-KO MuSC exhibit this feature of spontaneous activation by isolating single myofibers from adult extensor digitorum longus (EDL) muscles and quantifying projection length and number. Both projection length and number per single MuSC are significantly reduced in MuSC from TnC-KO mice compared to WT controls (**Figure 1F**). These findings indicate that TnC-KO MuSC exhibit a defect in quiescence in favor of premature myogenic commitment during postnatal muscle growth, leading to a decrease in the overall number of Pax7^+^ MuSC in TnC-KO adult skeletal muscles.

**Figure 1.**
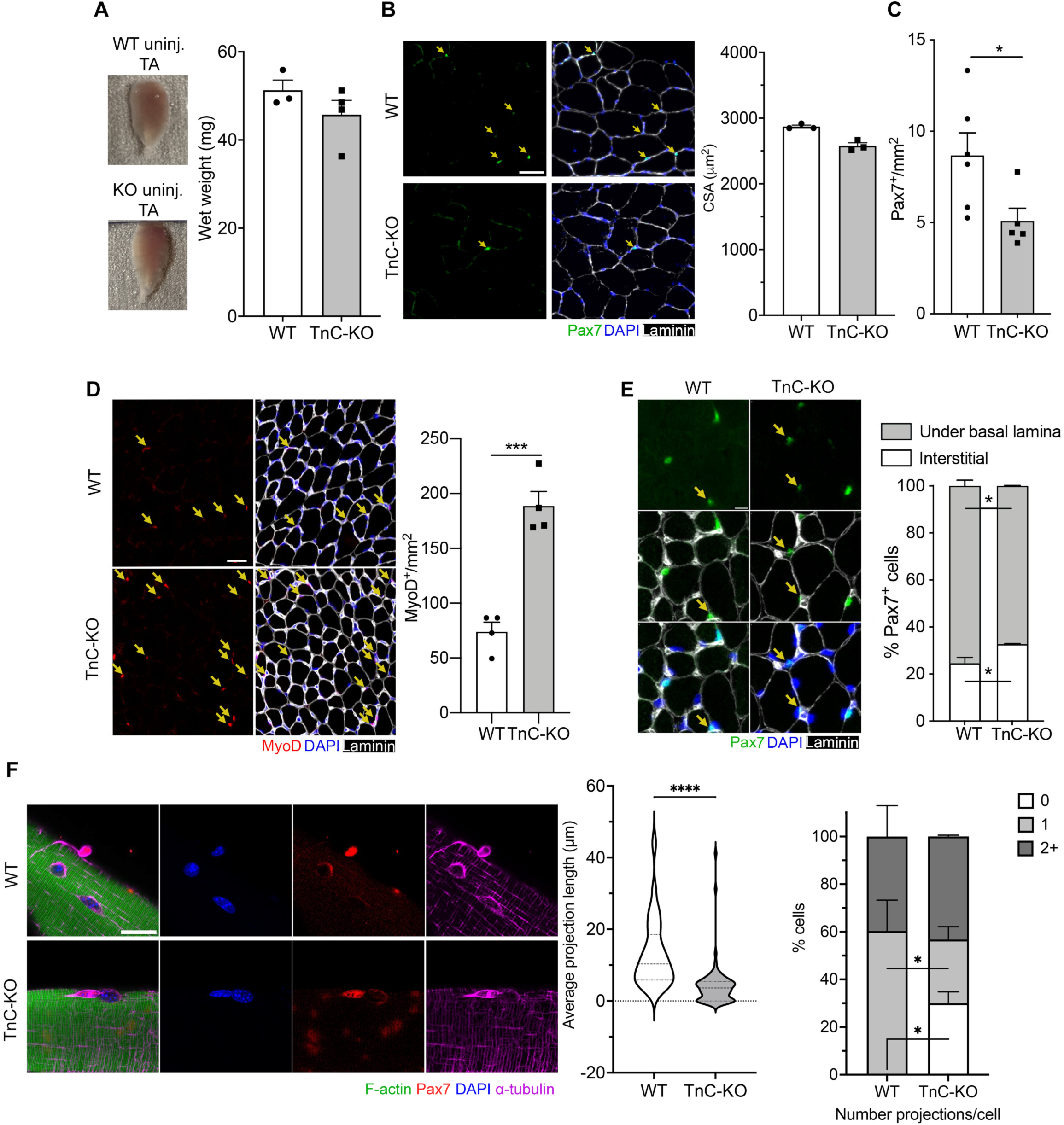
TnC is required for MuSC quiescence and pool maintenance. (A) Whole tibialis anterior (TA) muscles from wild type (WT) and TnC knockout (TnC-KO) adult (3-6 months old) mice and quantification of their wet weight (mg) (n≥3). (B) Representative immunofluorescence (IF) images of uninjured TA muscle cross-sections from WT and TnC-KO adult (3-4 months old) mice (Pax7, green; laminin, gray; DAPI, blue). Scale bar = 50 µm. Quantification of the cross-sectional area (CSA) in µm^2^ (n=3). (C) Quantification of the number of Pax7^+^ nuclei per mm^2^ (n≥5 mice). (D) IF images of cross-sections from P14 TA muscles of WT and TnC-KO adult (3-4 months old) mice (MyoD, red; laminin, gray; DAPI, blue). Scale bar= 50 µm. Quantification of the number of MyoD^+^ nuclei per mm^2^ (n=4). (E) Representative IF images of cross-sections from uninjured TA muscles of WT and TnC-KO P14 mice for quantification of Pax7^+^ cell localization (Pax7, green; laminin, gray; DAPI, blue). Scale bar= 10 µm. Quantification of Pax7^+^ cell localization under the basal lamina (quiescent) or in the interstitial space (activated) in WT versus TnC-KO muscles (n≥4). (**F**) Representative IF images of WT and TnC-KO freshly isolated myofibers from adult mice (F-actin, green; Pax7, red; ɑ-tubulin, magenta; DAPI, blue). Scale bar= 20 µm. Quantification of the distribution of quiescence projection length, and the number of projections per cell (n=3). Data are represented as mean ± SEM; \**p*<0.05, \*\*\**p*<0.001, \*\*\*\**p*<0.0001, *t* test (**A**, **B**, **C**, **D, E, F**); \**p*<0.05, Two-way ANOVA (**E, F**). Data are represented as the median with quartiles; \*\*\*\**p*<0.0001, *t test* (**F**).

### TnC is required for adult skeletal muscle regeneration and MuSC self-renewal

TnC is transiently expressed in regenerating TA muscles of adult WT mice following acute tissue injury induced by barium chloride injection ^18^. TnC expression begins to increase at 3 days post injury (DPI) and peaks at 5 DPI, as shown by both whole muscle Western blot analysis and histological analysis (**Figure 2A-B**). At 5 DPI, we observed a decreased cross-sectional area in TnC-KO muscles compared to WT controls, by assessing embryonic myosin heavy chain (eMyHC)-positive regenerating myofibers CSA (**Figure 2C**). Consistent with uninjured tissues, the number of Pax7^+^ MuSC was significantly lower in injured TnC-KO mice than WT controls at the same regeneration stage (**Figure 2D**). We also observed an increased number of MyoD^+^ committed progenitors at 5 DPI (**Figure 2E**), indicating increased myogenic commitment, as observed during postnatal muscle growth (**Figure 1D**). Histological analysis of adult TA muscles from TnC-KO and WT control mice confirmed that the reduction in MuSC number is maintained up to 30 DPI (**Figure 2F**). We then performed serial injury assays to investigate whether the absence of TnC impairs MuSC self-renewal. Injury was induced in TA muscles of both adult TnC-KO and WT control mice by barium chloride injection; one group was analyzed at 30 days after the first injury, while a second group was re-injured at 30 DPI in the same TA muscle and analyzed after an additional 30 days. Histology showed that the number of MuSC is further reduced after the second injury compared to the first injury in TnC-KO mice and to WT controls (**Figure 2F**). Overall, these findings indicate that the absence of TnC impairs adult MuSC self-renewal and leads to an inefficient skeletal muscle repair.

**Figure 2.**
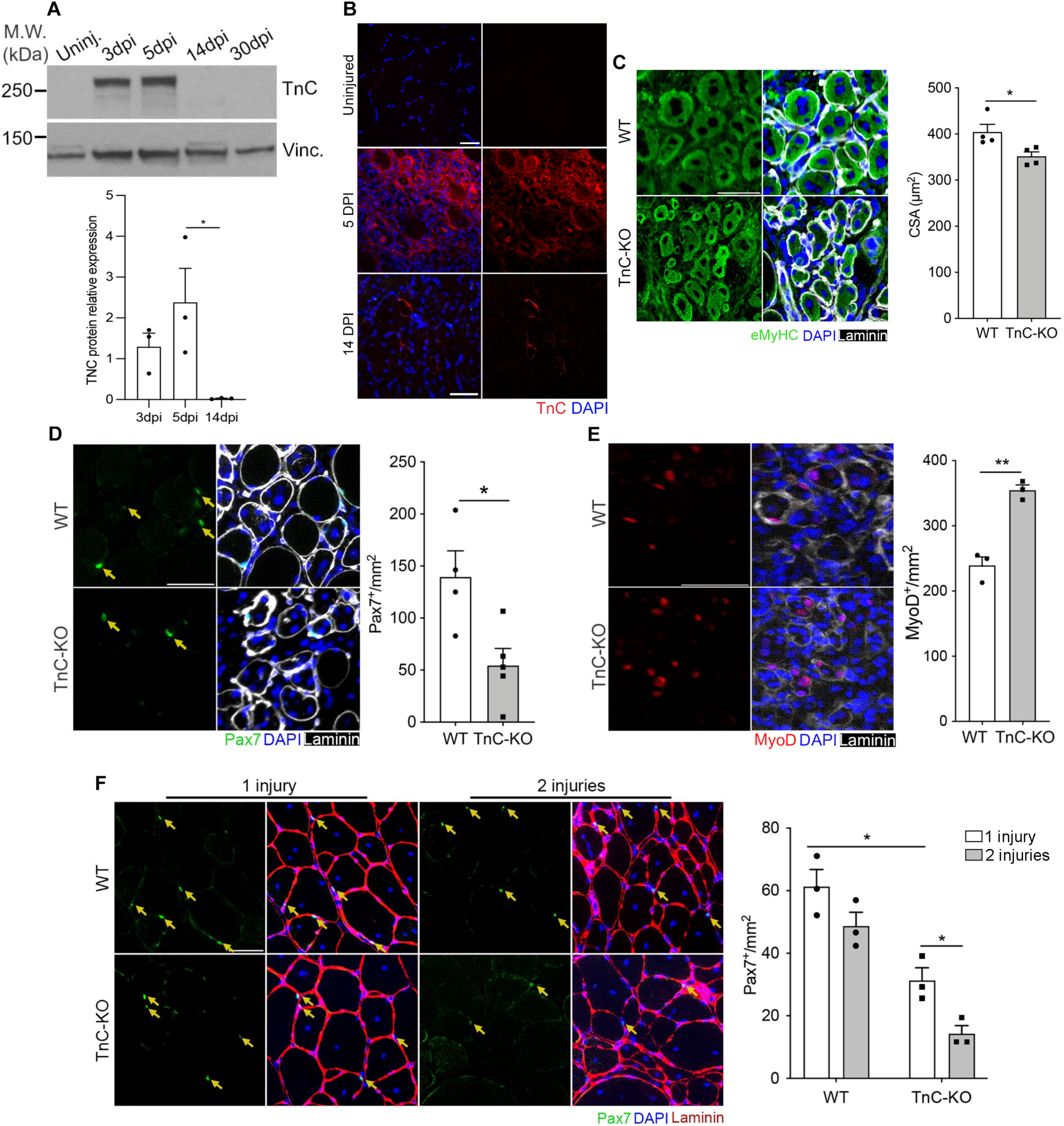
TnC expression peaks a 5DPI and promotes MuSC self-renewal. (A) Representative Western blot of TnC expression kinetics in uninjured and regenerating whole muscle protein lysates from adult (3-6 months old) mice and quantification (n=3). (B) Representative immunofluorescence (IF) images of cross-sections from uninjured and injured (5 DPI, and 14 DPI) TA muscles of WT adult mice (TnC red, DAPI blue). Scale bar= 50 µm. (C) Representative IF images for embryonic myosin heavy-chain (eMyHC) of injured TA muscles (5 DPI) in adult WT and TnC-KO mice (eMyHC, green; laminin, gray; DAPI, blue). Scale bar= 50 µm. Quantification of the cross-sectional area (CSA) (n=4). (D) Representative IF images of Pax7^+^ cells in injured (5 DPI) TA muscles in adult WT and TnC-KO mice (Pax7, green; laminin, gray; DAPI, blue). Scale bar= 50 µm. Quantification of the number of Pax7^+^ nuclei per mm^2^ (n=4). (E) Representative IF images of MyoD+ cells in injured (5 DPI) TA muscles in adult WT and TnC-KO mice (MyoD, red; laminin, gray; DAPI, blue). Scale bar= 50 µm. Quantification of MyoD^+^ cells per mm^2^ (n=3). (F) Representative IF images of Pax7^+^ cells in injured (30 DPI) or double-injured (30 DPI + 30 PDI) TA muscles in adult WT and TnC-KO mice (Pax7, green; laminin, red; DAPI, blue) (Scale bar= 50 µm) and quantification per mm^2^ (n=3). Data are represented as mean ± SEM; \**p*<0.05, one-way ANOVA (**A**); \**p*<0.05, \*\**p*<0.01, *t* test (**C**, **D, E**); \**p*<0.05, Two-way ANOVA (**F**).

### Fibro-adipogenic progenitors (FAPs) are the major source of TnC during skeletal muscle repair

To assess which specific cell types contribute to TnC deposition in skeletal muscle, we interrogated previously published single cell RNA-sequencing (scRNA-seq) datasets from mouse skeletal muscle at multiple time points after injury ^54^. scRNA-seq analysis at 5 DPI identified three muscle-resident cell types expressing TnC: FAPs (Pdgfra+/Ly6a+/Tnmd-), MuSC (Pax7+/Myod1+), and tenocytes (Tnmd+/Scx+) (**Figure 3A**), according to Oprescu *et al.* cell type classification. MuSC and FAPs exhibited a dynamic expression of the TnC transcript (**Figure 3B-C**), while expression of TnC in tenocytes remained high and did not change across uninjured and repairing muscles (**Figure S2A**). In MuSC, TnC expression was detected as soon as 0.5 DPI then declined back to uninjured levels by day 21 (**Figure 3B**). In FAPs, TnC expression began to increase at 0.5 DPI to be maintained at higher levels than MuSC, but declined by 10 DPI (**Figure 3C**). We validated these data by performing qPCR at the intermediate time points 3 and 7 DPI to confirm TnC expression during muscle repair in both MuSC and FAPs. (**Figure 3D-E**). The qPCR results confirmed overall higher levels of TnC mRNA in FAPs compared to MuSC, but with a decline of expression at an earlier timepoint than MuSC. To investigate cell-cell communication in the Tenascin signaling network between cell types in skeletal muscle, we utilized CellChat ^54-56^. In uninjured skeletal muscle, we identified tenocytes and FAPs as the highest probability senders of Tenascin signaling in uninjured skeletal muscle, along with lower probability of signaling from MuSC (**Figure 3F**). This analysis also recognized MuSC as the dominant receivers of Tenascin signaling in uninjured tissue along with milder contributions from tenocytes, FAPs, and immune cells (**Figure 3F, H**). FAPs overtake tenocytes to become the primary senders of Tenascin signaling at 5 DPI. We also observed a net increase in Tenascin signaling sent from MuSC and fibroblasts (**Figure 3G, I**). During repair, many more cell types are implicated as receivers of Tenascin signals; most notably this includes immune cells (M1, M2, Monocytes, and Proliferating Immune Cells), MuSC, FAPs, and tenocytes (**Figure 3G, I**). At 21 DPI, the sender-receiver dynamics returned to a state comparable with uninjured tissue (**Figure S2B-C**). Though Tenascin signaling is present in both uninjured and regenerating tissue, the relative contribution of specific ligands at each stage is unique. In uninjured tissue, a large part of the Tenascin signaling occurs through Tenascin-XB which shifts towards TnC at 5 DPI before returning primarily to Tenascin-XB at 21 DPI (data not shown). We speculate that there may be an increase in autocrine MuSC TnC signaling during muscle repair at 5 DPI that was not observed in the uninjured tissue or at 21 DPI (**Figure 3G, I** and **S2B-C**). Next, we performed transplantation assays to understand the function of TnC in MuSC and other cell types in the tissue microenvironment, and to characterize cell autonomous versus non-cell autonomous function in adult muscles. MuSC were isolated by FACS from hindlimb muscles of adult WT mice, and infected overnight with a lentivirus expressing red fluorescent protein RFP. After infection, 2,000 RFP^+^ MuSC were transplanted into the TA muscles of pre-irradiated adult TnC-KO or WT recipient mice, as previously described ^18^. At day 21 post transplantation, we observed significantly fewer donor-derived RFP^+^ myofibers in the TnC-KO host mice compared to WT hosts, indicating the functional relevance of TnC as a microenvironment-derived regulatory signal of MuSC-mediated myofiber formation (**Figure 3J**). We then investigated the role of FAP-produced TnC on MuSC behavior, due to the important role played by FAPs during skeletal muscle regeneration ^4-6^. MuSC and FAPs were isolated from adult (4-5 months old) WT and TnC-KO mice by FACS and co-cultured for 72 hours to assess the effect of cell-produced TnC to the maintenance of MuSC identity *in vitro*. Immunofluorescence analysis showed the presence of TnC in both WT MuSC and WT FAPs (**Figure 3K**), indicating that TnC production is preserved post-sort and after 72 hours in culture. We observed a greater number of Pax7^+^ cells when MuSC were co-cultured with WT FAPs, regardless of the MuSC genotype, when compared to WT MuSC monocultures (**Figure 3L**). However, there were fewer Pax7^+^ cells when MuSC were co-cultured with TnC-KO FAPs, which was comparable to what observed in WT MuSC monoculture (**Figure 3L**). These data suggest that FAP-derived TnC promotes maintenance of Pax7 expression in MuSC cultured *in vitro*. Together, these findings indicate that MuSC receive TnC signaling from cells in the tissue microenvironment, primarily FAPs and tenocytes, and that FAP-produced TnC regulates MuSC behavior *in vitro*.

**Figure 3.**
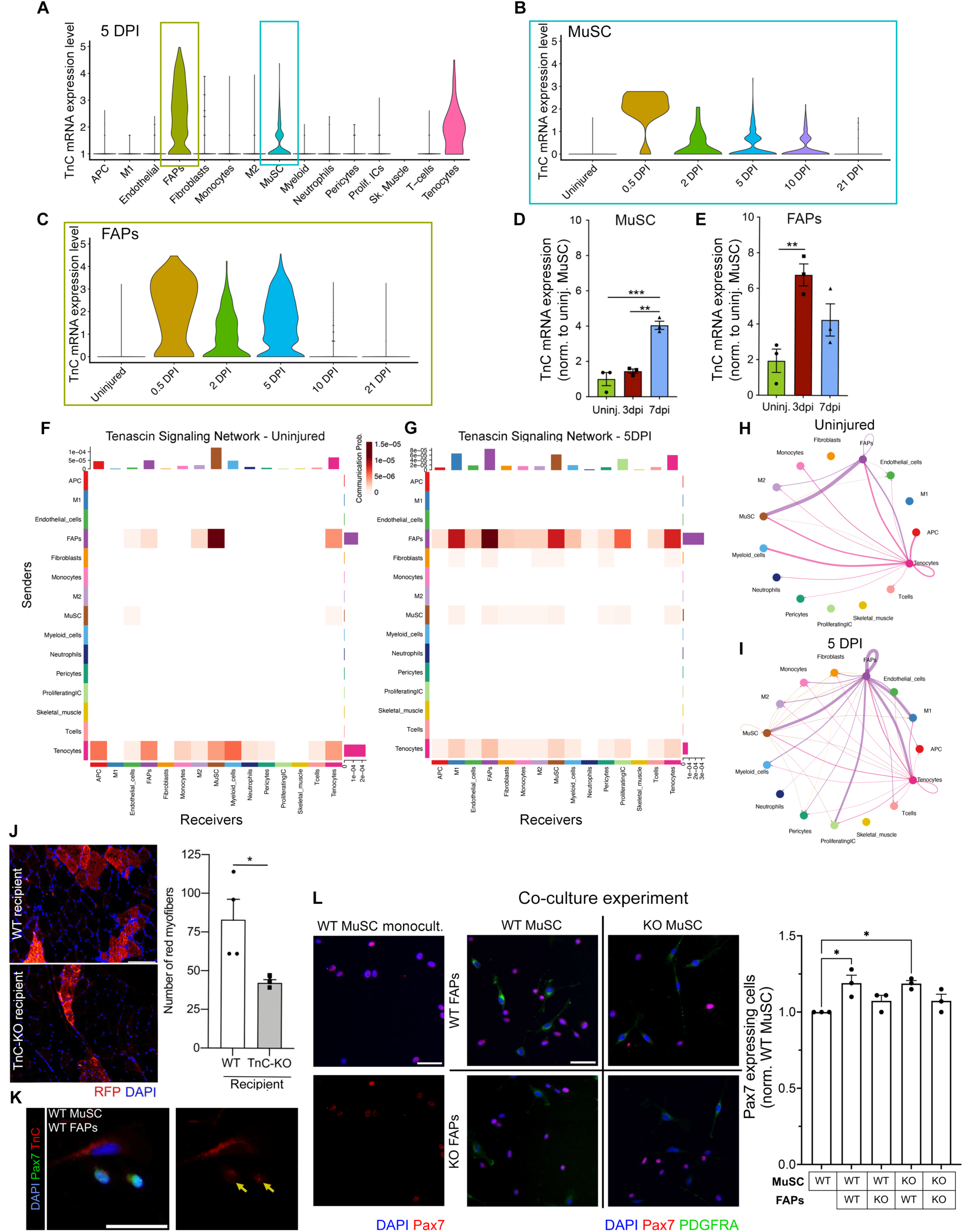
TnC from the tissue microenvironment is required for MuSC function. (A) scRNA-seq analysis of TnC expression at 5 DPI TA muscles of adult WT mice. Data originally from Oprescu *et al.*, 2020. (**B**, **C**) TnC mRNA expression dynamics in MuSC and FAPs from uninjured and regenerating TA muscles at different DPIs (0.5 – 21). Data originally from Oprescu *et al.*, 2020. (**D, E**) TnC mRNA expression dynamics in MuSC and FAPs in uninjured and at 3 and 7 DPIs with qPCR. (**F, G**) Sender-Receiver Probability Heatmap of the tenascin signaling network in uninjured and 5 DPI TA muscles. Data originally from Oprescu *et al.*, 2020. (**H, I**) Chord diagrams representing incoming signaling to MuSC in uninjured and 5 DPI TA muscles. Data originally from Oprescu *et al.*, 2020. (**J**) IF images representing the contribution of transplanted RFP-labeled WT MuSC from adult 3 months old donor mice to regenerating TA muscles of WT or TnC-KO age-matched recipient mice (regenerating fibers – RFP, red, DAPI blue) (Scale bar= 50 µm) and quantification of RFP+ myofibers (ný3 mice). (**K**) Representative IF images of WT MuSC/WT FAP co-cultures from adult mice (Pax7, green; TnC, red; DAPI, blue). Yellow arrows indicate TnC in MuSC. Scale bar= 50 µm. (**L**) Representative IF images of WT MuSC monocultures (Pax7, red; DAPI blue) (scale bar= 50 µm) and co-cultures of MuSC and FAPs from WT and TnC-KO adult mice (Pax7, green; TnC, red; DAPI, blue) (Scale bar= 50 µm). Quantification of Pax7+ cells normalized on WT MuSC monoculture. Data are represented as mean ± SEM; \*\**p*<0.01, \*\*\**p*<0.001, one-way ANOVA (**D, E**), \**p*<0.05, *t* test (**J**), \**p*<0.05, one-way ANOVA (**L**).

### TnC signals through surface protein Annexin A2 to regulate MuSC function

Although previous studies have described several cell surface receptors mediating TnC signaling, including Annexin A2 (Anxa2), Syndecan-4, TLR4, EGFR, integrin ⍺5/β1, ⍺8/β1 and ⍺9/β1 in multiple cell types ^35^, these interactions are less well understood in the context of skeletal muscle regeneration. mRNA expression by scRNAseq analysis in MuSC for these known TnC receptors, both before injury and during regeneration, revealed that ⍺5/8/9 integrin transcripts were not detected. While syndecan-4 progressively decreased during MuSC activation, TLR4 and EGFR mRNA levels were not notably changed across the repair process (data not shown). We analyzed the dynamics of RNA expression at the single cell level in MuSC and FAP populations for the key target transcripts TnC and Anxa2, and cell-identifying transcription markers (Pax7 for MuSC, Pdgfra for FAPs). We found that Anxa2 expression transiently increases in MuSC at 3.5 and 5 DPI (**Figure 4A**). In contrast, there is no evident change to the expression of Anxa2 in FAPs throughout the muscle regeneration process, except for a minor decrease at 21 DPI (**Figure 4A**). These data also show that TnC expression begins earlier in FAPs than MuSC, and that FAPs are greater producers of TnC than MuSC during regeneration (**Figure S3A**). Next, we detected the physical interaction of TnC with Annexin A2 from 5 DPI adult (3-5 months old) TA muscles through co-immunoprecipitation assays (**Figure 4B)**. Immunofluorescence analysis on freshly isolated MuSC confirmed that Annexin A2 is expressed in Pax7^+^ MuSC after 24 hours in culture (**Figure 4C**). We then looked at what percentage of AnxA2^+^ or AnxA2^-^ MuSC (Pax7^+^) or FAPs (Pdgfra^+^) express TnC and the relative expression of TnC in these populations throughout the regeneration process. The observed mRNA expression patterns for each cell type indicate that not only is the average expression of TnC increasing per cell, but also that a greater proportion of Anxa2^+^ cells transiently express TnC during regeneration in both MuSC and FAP populations (**Figure 4D**). We also investigated the Anxa2^-^ fraction of MuSC and FAPs to evaluate whether these different subpopulations underwent unique dynamics in TnC expression. Interestingly, we observed a reduced number of analyzed cells (raw data of cell numbers) per condition overall in the AnxA2^-^ compared to the AnxA2^+^ cell fractions (**Figure S3B**), and most importantly that Anxa2^-^ MuSC express low levels of TnC during regeneration, while Anxa2^-^ FAPs follow a similar, but reduced trend in comparison to their Anxa2^+^ counterparts (**Figure 4D**). A weaker trend was also observed in committed Myod1^+^/Anxa2^+^ MuSC, with reduced raw cell count compared to the corresponding Pax7^+^ populations (**Figure S3C**). These data suggests that the subpopulation of Anxa2^+^ cells are responsible for driving the majority of TnC signaling in both MuSC and FAPs. To assess the requirement of Annexin A2 to mediate the effects of TnC on MuSC, we performed a knockdown by infecting freshly isolated mouse MuSC with lentivirus carrying shRNA for Annexin A2. After infection (24 hours), MuSC were further cultured in growth conditions either in the presence or absence of recombinant TnC protein for 48 hours. The efficiency of knockdown for Annexin A2 was >75% at 48 hours from infection (**Figure S3D**). In these conditions, we observed that the treatment with TnC promoted an increase in the number of Pax7^+^ cells compared to non-treated GFP-control MuSC, while knockdown of Annexin A2 abrogated the effect of TnC (**Figure 4E**). Additionally, the TnC treatment showed a specific effect on Pax7^+^ MuSC, while leaving the numbers of MyoD^+^ myogenic progenitors unaffected in both control and knockdowns (**Figure S3E**). Therefore, these results indicate that Annexin A2 expressed on MuSC is required for TnC signaling to occur in this context.

**Figure 4.**
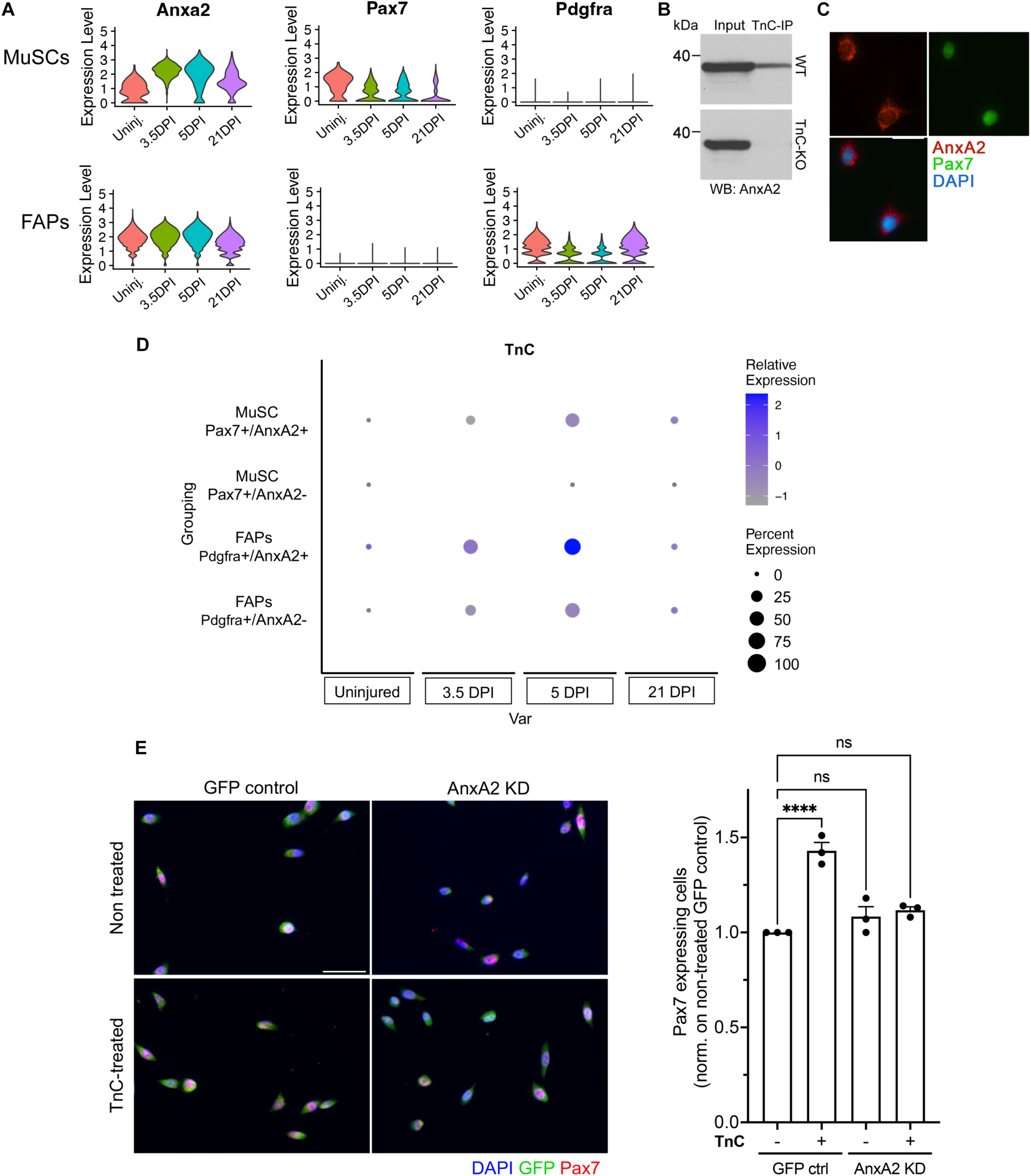
TnC signals through Annexin A2 to maintain MuSC in quiescence. (A) Violin plots depicting the different expression dynamics AnxA2, Pax7, and Pdgfra at different DPIs (0, 3.5, 5, 21) in MuSC and FAPs (online available dataset). (B) Western blot analysis for Annexin A2 on TnC-immunoprecipitated 5 DPI TA muscle protein lysate samples. (C) Validation by immunofluorescence (IF) of Annexin A2 expression in WT MuSC *in vitro* (Pax7, green; Annexin A2, red; DAPI, blue) (Scale bar= 20µm). (D) Dot plot representing the changes in the relative expression of TnC and the percentages of TnC expressing cell populations within Pax7^+^/AnxA2^+^ and Pax7^+^/AnxA2^-^ MuSC and Pdgfra^+^/AnxA2^+^ and Pdgfra^+^/AnxA2^-^ FAPs in uninjured TA muscles and across different timepoints (3.5, 5, 21 DPI). Data originally from Oprescu *et al.,* 2020. (E) Representative IF images of GFP control and AnxA2 knockdown (KD) being treated or not with recombinant TnC (48h treatment) (GFP, green; Pax7, red; DAPI, blue) (scale bar= 50 µm) and quantification of Pax7^+^ cells normalized on non-treated GFP control MuSC (n=3, N_fov_=30). Data are represented as mean ± SEM; \*\*\*\**p*<0.0001, one-way ANOVA (**E**).

### TnC promotes MuSC migration

The ability of MuSC to migrate upon injury is an essential feature that allows these cells to reach the damaged area and initiate the regeneration process. TnC has been previously shown to promote migration in multiple cell types including fibroblasts, endothelial and smooth muscle cells, astrocytes, and in metastatic cancer ^57-59^. Thus, we asked whether TnC regulates MuSC ability to migrate by performing time-lapse microscopy on freshly isolated MuSC from TnC-KO and WT mice cultured in growth conditions for 24 hours. Both the total traveled distance and the average velocity of the cells were significantly reduced in TnC-KO MuSC compared to WT controls (**Figure 5A-C**). We implemented transwell assays to further assess MuSC migration capacity; we observed that the lack of TnC reduced MuSC migration after 48 hours in culture (**Figure 5D**). We confirmed that the increased number of cells migrated through the transwell was indeed due to migration and not to MuSC proliferation (**Figure 5E**). Next, we administered soluble recombinant TnC to WT MuSC, which further promoted MuSC migration compared to untreated WT cells in transwell assays (**Figure 5F**). Consistent with the *in vivo* data, MuSC proliferation was not altered by the presence of soluble recombinant TnC (**Figure 5F**). We further assessed cytoskeleton organization of the myofibers from TnC-KO mice, to explore whether TnC gene deletion affects their structure. Immunofluorescence analysis of ɑ-tubulin revealed that the cytoskeletal network of freshly isolated myofibers from TnC-KO mice was more disorganized than that of WT control myofibers, (**Figure S4A**). We observed a large dispersion spread in the directionality of microtubules in the TnC-KO, compared to the WT control, which conversely showed a very narrow range or dispersion (**Figure S4B-C**). Together, these findings indicate that the deletion of TnC simultaneously limits MuSC migration *in vitro* and causes disorganization of the myofiber microtubular network.

**Figure 5.**
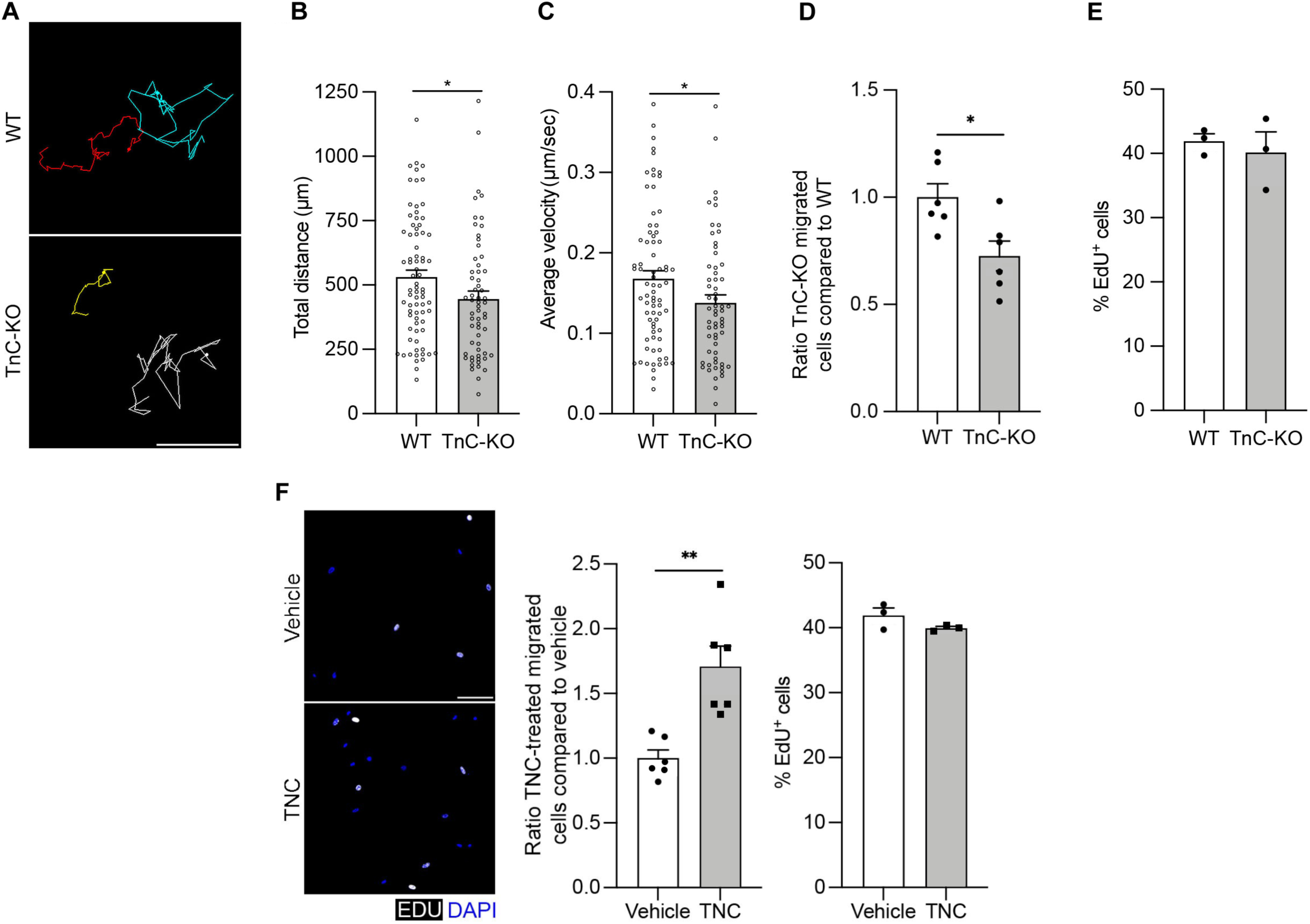
TnC promotes MuSC migration *in vitro*. **(A)** Representative images of time-lapse microscopy to track cultured MuSC from adult WT and TnC-KO mice. Scale bar= 50 µm. **(B)** Quantification of total distance covered (n= 3, N_cell_≥75). **(C)** Quantification of average velocity of cells (n= 3, N_cell_≥75). **(D)** Quantification of migrated TnC-KO cells through transwell matrix normalized on WT (n=6). **(E)** Quantification of the percentage of proliferating cells through EdU staining (timepoint= 48 hours) (n=3). **(F)** Representative immunofluorescence (IF) images of migrated WT MuSC from adult (3-5 months old) mice with or without recombinant TnC treatment (EdU gray, DAPI blue) (scale bar= 50 µm). Quantification of normalized migrated TnC-KO cells through transwell matrix after TnC treatment (timepoint= 48 hours) (n=6) and quantification of the percentage of proliferating cells through EdU staining (timepoint= 48 hours) (n=3). Data are represented as mean ± SEM; \**p*<0.05 and \*\**p*<0.01, *t* test.

### TnC KO mice exhibit a premature aging phenotype in skeletal muscle

In light of the role played by TnC during muscle regeneration and the widely studied impairment of regeneration in aged skeletal muscle, we asked whether TnC expression changes during skeletal muscle aging. Comparison of young (3-5 months old) and aged (18-26 months old) regenerating muscles demonstrated decreased levels of TnC both via Western blot analysis (**Figure 6A**) and immunofluorescence (**Figure 6B**). We also showed that the number of MuSC was reduced in aged muscles compared to young controls (**Figures 6C** and **S5A**), consistent with previous literature ^60-62^. TnC deletion significantly reduced the wet weight of uninjured TA muscles from aged (23-29 months old) TnC-KO mice compared to both young and aged WT TA controls (**Figure S5B**). We also recorded reduced fibrosis in aged TnC-KO TA muscle cross-sections compared to young and aged WT and young TnC-KO samples (**Figure S5C**), but did not observe any accumulation of adipose tissue across genotypes and ages (data not shown). We further observed reduced myofiber CSA in uninjured aged TnC-KO mice compared to each of WT young, TnC-KO young, and WT aged mice (**Figure 6D**). A comparable reduction in CSA was also observed in the diaphragm (**Figure S5D**), indicating that the observed phenotype is not limited to skeletal muscles of the hindlimb. We also detected a significantly decrease in the number of Pax7^+^ cells in aged TnC-KO mice compared to the age-matched WT counterparts (**Figure 6E**). Following tissue injury by barium chloride injection, we observed impaired tissue repair in aged WT muscles compared to young controls. At 5 DPI, the damaged areas in old TA contained a reduced area of eMYHC^+^ fibers, fewer regenerating myofibers, and smaller regenerating fiber CSA as compared to young (**Figure 6F** and **S5E**). Young TnC-KO muscles did not contain a significantly reduced number of regenerating myofibers following injury (**Figure S5E**), while both the area of muscle comprised of eMYHC^+^ fibers as well as the number of eMyHC+ myofibers were lower than in Young WT muscles and were comparable to aged WT controls (**Figure 6F** and **S5E**). Aged TnC-KO muscles undergoing regeneration showed an even greater decline in the percentage of eMYHC^+^ area and in regenerating fiber CSA compared to both aged WT and young TnC-KO muscles (**Figure 6F** and **S5E**). We then used transwell assays and MuSC migration as a readout to further investigate this regeneration phenotype in aged muscle; we observed that aged MuSC exhibited a defect in cell migration compared to young controls (**Figure 6G**). When aged MuSC were exposed to TnC in the transwell assay, the treatment rescued their migratory defect (**Figure 6H**). Together, these data demonstrate that the lack of TnC in aged mice further impairs the regeneration phenotype observed in young TnC-KO mice, and that the impaired migration observed in aged muscle caused by the decreased levels of TnC is rescued by treating aged MuSC with exogenous TnC.

**Figure 6.**
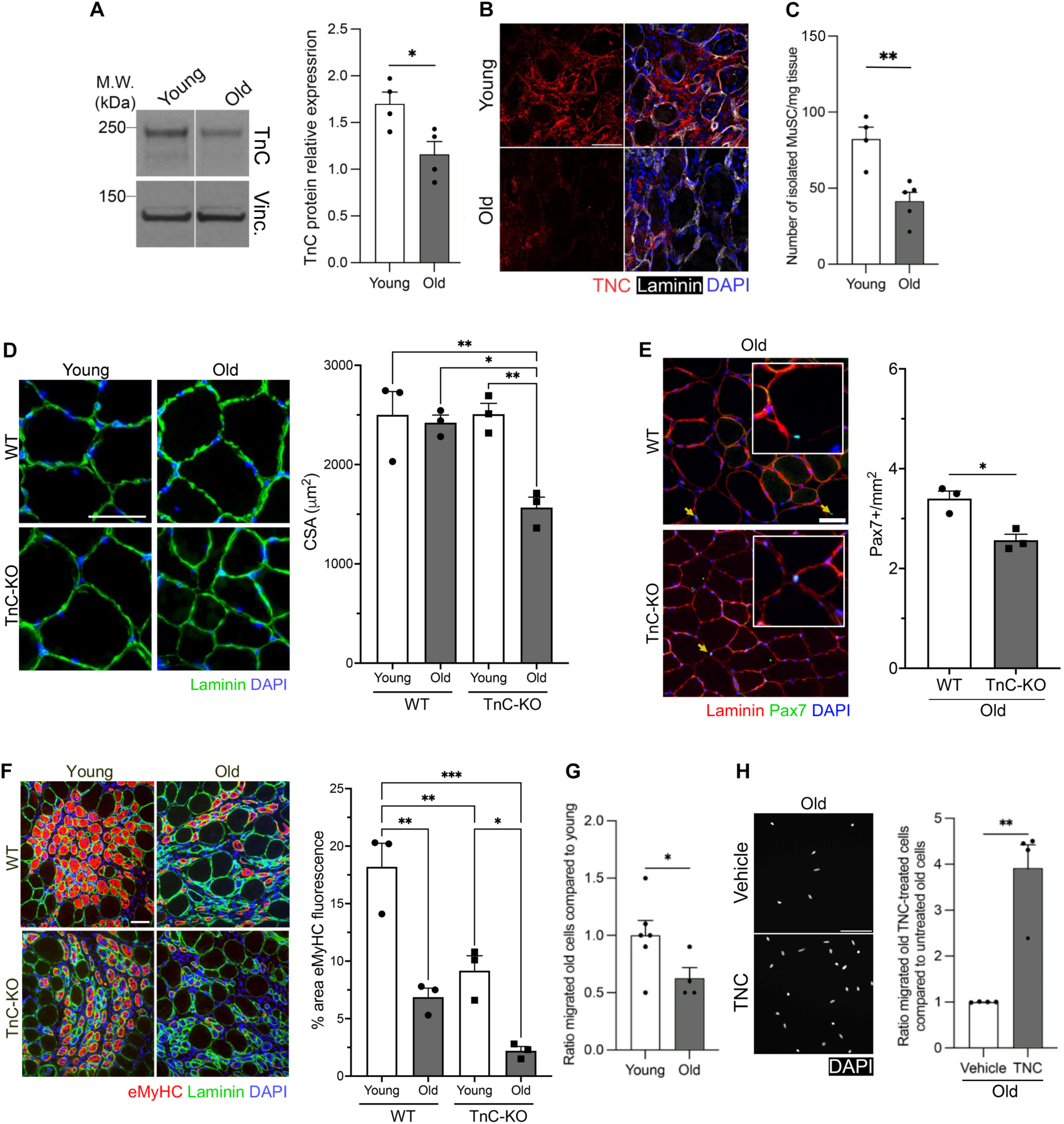
TnC declines with aging and is required for muscle maintenance and regeneration of aged skeletal muscles. (A) Western blot for TnC on young (3-5 months old) and old (18-26 months old) WT mice and quantification of TnC protein relative expression (n=4). (B) Representative immunofluorescence (IF) images of 5 DPI TA muscles of young and old mice (TnC red; laminin gray; DAPI blue). Scale bar= 50 µm. (C) Quantification of the number of FACS isolated MuSC per mg of muscle tissue in young and old mice (ný4 mice). (D) Representative IF images of cross-sections of uninjured TA muscles from young (4-6 months old) and old WT (23-29 months old) and TnC-KO mice (laminin, green; DAPI, blue) (scale bar= 50 µm) and quantification of the cross-sectional area (CSA) in µm^2^ (n=3). (E) Representative IF images of cross-sections of uninjured TA muscles from old WT and TnC-KO mice (Pax7, green; laminin, red; DAPI blue) (scale bar= 50 µm) for Pax7^+^ cell quantification (n=3). (F) Representative IF images of cross-sections of injured (5 DPI) TA muscles from young and old WT and TnC-KO mice (laminin, green; eMyHC, red; DAPI, blue) (scale bar= 50 µm) and quantification of the percentage of regenerating area (eMyHC^+^) (n=3). **(G)** Quantification of migrated aged WT MuSC through transwell matrix (48 hours culture) normalized on WT young mice (n≥4). **(H)** Representative IF images of migrated cells with or without recombinant TnC treatment (timepoint= 48 hours) (DAPI, gray) (scale bar= 50 µm) and quantification of migrated aged WT MuSC through transwell matrix after TnC treatment normalized on control non-treated cells (n=4 mice). Data are represented as mean ± SEM; \**p*<0.05, \*\**p*<0.01, *t* test (**A, C, E, G, H**), \**p*<0.05, \*\**p*<0.005, \*\*\**p*<0.0001, one-way ANOVA (**D, F**).

In conclusion, there is a consistent phenotype of TnC-KO mice, where young muscles have fewer MuSC, a reduced myofiber CSA, impaired migration, and a decreased ability to repair tissues following injury. We also showed that exposing TnC-KO mice to soluble recombinant TnC protein rescued the migratory capabilities of TnC-KO MuSC.

## DISCUSSION

### TnC promotes MuSC self-renewal and migration

During mouse postnatal muscle growth, MuSC enter a quiescence state around one month of age ^18,48-51^. Our findings show that MuSC numbers are not affected during prenatal development in TnC-KO mice, but that deletion of TnC reduced the number of MuSC by one month of age. We also provide evidence that TnC-KO mice have increased localization of MuSC outside of the sublaminar niche and are located in the interstitial space. In addition, in TnC-KO we observed a higher number of cells expressing MyoD, a marker of committed progenitors, compared to WT controls. These findings suggest that MuSC undergo spontaneous myogenic commitment and fail to enter or maintain quiescence in the absence of TnC. Recent studies using intravital imaging have demonstrated that a morphological feature of quiescent MuSC is the presence of long cellular projections ^63-65^. Upon tissue injury, rearrangement of cytoskeletal components occurs, and the retraction of these long cellular projections is observed upon MuSC activation. This is mediated by mechano-sensing pathways triggered by a Rac-to-Rho switch and Piezo-1 signaling ^52,53^. Our data demonstrate that TnC-KO MuSC exhibit both reduced length and number of cellular projections in the absence of TnC, which further supports spontaneous MuSC activation. A potential role of MuSC projections is to probe their immediate surrounding tissue for myofiber damage and respond by retractions and MuSC activation. In our study, we have found that TnC-KO myofibers show a disorganized cytoskeleton, which may act as a mechanical stress for MuSC and thus triggers activation even under steady-state conditions. To address the question of whether this phenotype is associated with a defect in MuSC self-renewal, we performed serial injury assays. Upon repeated injuries, we observed fewer MuSC *in vivo* indicating that self-renewal is impaired in TnC-KO mice. TnC could be a regulator of the division dynamics of MuSC and play a role balancing symmetric and asymmetric divisions. Our previous work has shown that fetal MuSC, which express TnC, undergo asymmetric division and symmetric expansion in culture, compared to adult MuSC which mainly sustain symmetric depletion events ^18^. We propose that the absence of TnC biases MuSC towards symmetric depletion events, resulting in defective self-renewal, maintenance, and re-entry into quiescence following tissue injury. We further show that MuSC isolated from TnC-KO mice exhibit impaired migration, both by time-lapse microscopy and in transwell assays. Administration of soluble TnC to WT MuSC enhanced their migration. TnC has previously been reported to promote migration in multiple tissues and cell types by inhibiting focal adhesion formation and regulating cytoskeleton organization ^38,66-68^. Whether the impaired migration in the absence of TnC is a cause or consequence of an activated MuSC phenotype is currently unclear, and future studies will enable to shed light on this functional interdependence. We suggest that TnC plays a role in migration through the engagement of MuSC receptors that are upregulated at the cell surface during regeneration.

### TnC engages Annexin A2 to promote MuSC maintenance and expansion

We demonstrate that TnC signaling in MuSC occurs through binding to Annexin A2. TnC has been previously reported to bind several cell surface receptors, including Annexin A2, Syndecan-4, EGFR, TLR4, as well as integrins ^35^. Our analysis of skeletal muscle scRNAseq available data showed that the transcripts for Syndecan-4, EGFR, TLR4, and the aforementioned integrins are either not detected or do not significantly change in MuSC during the regeneration process. Although we cannot rule out the role of these surface receptors in the context of skeletal muscle repair, we focused on Annexin A2, as this surface molecule demonstrated the greatest changes in mRNA expression during the early stages of regeneration, following an expression trend similar to that of TnC.

The role of Annexin A2 has been previously studied in the context of myofiber membrane repair ^39,69^. Annexins are a widely expressed family of Ca^2+^- and phospholipid-binding proteins that contribute to many cell functions at the plasma membrane including membrane organization and trafficking, transmembrane channel activity, cell-ECM communication, and often act as a bridge for membrane-cytoskeleton interactions ^70^. In skeletal muscle, Annexin A2 trafficking to the injured site ^71^ and its interaction with proteins associated with sarcolemma repair, such as dysferlin ^72-74^ are required processes for efficient muscle regeneration. Indeed, myofibers lacking Annexin A2 demonstrated defective repair upon muscle injury ^73^. Annexin A2 is upregulated in intestinal epithelial cells during migration and promotes cell spreading and positively regulates Rho activity ^75^. Upon interaction with TnC, Annexin A2 promotes the loss of focal adhesions and cell migration in multiple cell types ^38,76^.

Our bioinformatics analysis shows that upon tissue injury Annexin A2 transiently increases in expression to then return to uninjured levels at 21 DPI. We observed that TnC-KO MuSC exhibited defects in activation and migration, which is in line with previous evidence that Annexin A2 mediates actin rearrangements through Rho activation ^77^. We propose that the transient increase of MuSC-specific Annexin A2 expression during tissue repair might promote TnC-mediated MuSC activation and migration to the damaged area during the early stages of regeneration. This may occur through a feedback loop involving the reciprocal regulation of TnC:Annexin A2 interaction and the Rho/ROCK axis. Our expression analysis has highlighted that not all MuSC exhibit the same level of expression of Annexin A2. Whether these differences underlie distinct cell subpopulations or transitional cellular states it is currently unclear, and it will be interesting to address it in future studies.

### FAPs as cellular sources of TnC during muscle regeneration

We have previously shown that TnC is expressed by MuSC at fetal stages as well as in adult MuSC during tissue repair ^18^. Downregulation of TnC in fetal MuSC, though not in adult MuSC, impairs MuSC contribution to skeletal muscle repair upon transplantation. This different behavior suggests that multiple cell types in the adult tissue microenvironment produce TnC to achieve efficient regeneration ^18,23^. Our transplantation assays demonstrate that when WT adult MuSC are transplanted into TnC-KO mice, their contribution to tissue repair is impaired, indicating that TnC from the tissue microenvironment is required for MuSC function in this context. Our analyses of scRNAseq and qPCR data demonstrate that the major sources of TnC in skeletal muscle during tissue repair are FAPs, tenocytes, and MuSC. TnC has a known anatomical localization at the tendons ^19,78^. TnC transcript levels are indeed high and constant in tenocytes during both homeostasis and repair, thus we speculate that these cells do not contribute strongly during regeneration. Conversely, FAPs dynamically increase expression of TnC in the initial phases of tissue repair. Our co-culture experiments demonstrate that TnC released from FAPs is required for the maintenance of the MuSC pool. When FAPs from TnC-KO mice are co-cultured with WT MuSC, we did not observe the expansion of the Pax7^+^ population *in vitro*, indicating that FAP-derived TnC is required in this process. While it has been previously shown that co-cultivating FAPs and MuSC isolated from uninjured mouse muscle for 7-10 days increases the terminal differentiation of myogenic progenitors ^4,5,79,80^, our results indicate that upregulation of TnC in FAPs occurs specifically at early timepoints, suggesting that FAPs might promote different processes at different stages of during tissue repair. Our bioinformatic analysis of scRNAseq data further shows that 50-75% of FAPs upregulate TnC in response to injury, demonstrating the heterogeneity of this cell population in this context. It is currently unclear whether these differences underlie distinct cell subpopulations or transitional cellular states. During wound repair, TnC upregulation is induced by TGFb signaling mediated by with SP1 or ETS1 in a complex with CBP/p300 in fibroblasts ^81^. In addition, TnC gene transcription has been shown to be induced by mechanical stress through ⍺5β1 integrin, ILK, and Rho/ROCK signaling, inducing nuclear translocation of MLK1 (MRTF-A) ^82,83^. We suggest a novel role for FAPs in regeneration through the secretion of TnC, where initially FAP-derived TnC is required to regulate the number of MuSC following injury, that is then followed by the known role of FAPs to induce differentiation of activated MuSC.

### Aged skeletal muscle shows decreased levels of TnC

Our results show that TnC expression during tissue repair is reduced during aging. We further show that mice lacking TnC exhibit a premature aging phenotype, with a reduced number of MuSC and impaired MuSC self-renewal and overall tissue regeneration. Analysis of TnC-KO muscles upon injury shows that the defect is further exacerbated during aging, with a severe reduction in the number of MuSC and severely impaired tissue repair. Previous studies have reported an increased abundance of FAPs in aged muscles, while other studies show a decrease in the number of FAPs in favor of differentiation towards a fibrotic state ^45,46,84^. We observed neither increased fibrosis nor adipogenesis in our aged TnC-KO muscles, suggesting that the lack of TnC in aged muscle does not bias FAPs towards either cellular fate. However, the reduced levels of TnC in aged WT muscles could be the result of either changes in TnC expression or changes in FAP abundance.

Delivering an ECM protein as large as TnC *in vivo* is inherently challenging as it will mostly accumulate at the site of injection and not distribute across tissues evenly. However, recent studies have found TnC localized within extracellular vesicles (EVs) secreted from fibroblasts and breast cancer cell lines ^85,86^, and we have detected TnC protein in EVs from skeletal muscle FAPs (data not shown). Thus, we envision that future approaches leveraging EVs as the delivery system for TnC would provide a potential therapeutic approach to restore signaling and promote tissue repair in aged muscles.

### Limitation of the study

Understanding the cellular and molecular complexity of a regenerating tissue is a current and challenging question. This study provides novel insights on the role of the tissue microenvironment in regulating MuSC function during muscle repair, with particular focus on a novel role of FAPs and ECM protein TnC. Our knockdown experiments and co-culture assays have demonstrated the contribution of FAP-derived TnC in MuSC function, however one limitation of this study is the lack of inducible FAP-specific TnC knockout or MuSC-specific Annexin A2 knockout mice, currently not available. These models will enable future studies to investigate the role of TnC:Annexin A2 interactions *in vivo*. This study did not incorporate the assessment of TnC epigenetic regulation and post-translational modifications (PTMs). Therefore, future epigenetic approaches and proteomic studies will identify the changes in regulatory elements and PTMs during regeneration versus homeostasis, and in the context of aging. While we have investigated the migration and activation phenotype in the context of TnC absence, both *in vitro* and *in vivo*, an exciting question lays in the timing and relationship between these two processes. Genetic tools would need to be developed to address this biological challenge. Finally, while we plan to lead future studies to deliver TnC *in vivo* to promote muscle regeneration, our treatments with TnC *in vitro* have already shown the potential of TnC as a therapeutic tool, through its effect in promoting cell migration and cell fate decisions.

## ACKNOWLEDGMENTS

This work was supported by the Association Française contre les Myopathies (AFM 21080), and the US National Institute of Health (NIH) grants R01 AR064873, R01 AR077448 and R21 AR075205 to AS. We thank the members of the Puri and Colas labs, in particular Drs. L. Caputo and X. Wei for their immense support for qPCR and FACS experiments. We thank Drs. Faessler and Orend for the TnC-KO mice. We thank the researchers who made their datasets publicly accessible, especially Dr. Shiuhan Kuang, for carrying out respectively the scRNA-seq, which was implemented in this study. We thank the following people at the SBP Core Facilities for technical support: B. Charbono, T. Omel, D. Sandoval and A. Vasquez from the Animal Facility; Y. Altman, A. Cortez, and B. Portillo from the Flow Cytometry Core Facility; L. Boyd from the Cell Imaging Facility. G. Garcia and M. Sevilla from the Histology Core Facility. We apologize to authors whose papers we could not cite due to space limitations.

## AUTHOR CONTRIBUTIONS

Conceptualization, M.L., A.C., and A.S.; Methodology, M.L., A.C., C.K., C.S., A.R., C.N., A.K., G.G., and D.M.; Investigation, M.L., A.C., C.K., C.S., G.G., and A.S.; Writing & Editing, M.L., A.C., C.K., C.S., A.R., C.N., A.K., G.G., D.M., A.Co., P.L.P., and A.S.; Funding Acquisition, A.S.; Resources, A.S.; Supervision, A.S.

## COMPETING FINANCIAL INTERESTS

The authors declare no competing financial interests.

## METHODS

### Contact for reagent and resource sharing

Further information and requests for resources and reagents should be directed to and will be fulfilled by the Lead Contact, Alessandra Sacco, Ph.D.

### Experimental model and subject detail

#### Experimental Animals

Tenascin-C knock out (TnC-KO) mice were a kind gift from Drs. Faessler and Orend. All protocols were approved by the Sanford Burnham Prebys Medical Discovery Institute Animal Care and Use Committee. Mice were housed according to institutional guidelines, in a controlled environment at a temperature of 22°C±1°C, under a 12-hour dark-light period and provided with standard chow diet and water ad libitum. Male and female TnC-KO and wild-type mice (FVB background) at ages E16.5, P14, P30, adult age (3-6 months old), and geriatric (18-29 months old) were used. All mice were maintained in FVB background. Adult NOD/SCID were used for the transplantation experiments.

### Method details

#### Cells Isolation

Muscle stem cells (MuSC) were isolated as describes in Gromova et al., 2015 with minor revisions ^87^. Whole hindlimb (tibialis anterior, extensor digitorum longus, gastrocnemius, soleus, plantaris, vastus lateralis, quadriceps, and adductors) were minced and sequentially incubated in 800 units/mL collagenase type II solution (catalog number: 17101-015, Life technologies, Gibco^®^) and subsequent incubation with 80 units/mL collagenase II and 2 units/mL dispase II (catalog number: 04942078001, Roche) solution. Muscle tissues were then passed through a 10 mL syringe with 20 G needle and a 40 μm nylon filter. Primary and secondary antibodies incubation was performed in a 300 µL volume. Biotin-labeled lineage negative cells (CD45^+^, CD11b^+^, CD31^+^, Sca1^+^ cells) were either depleted using streptavidin beads (catalog number: 130-048-101, Miltenyi Biotec) through magnetic field or excluded during sorting by using streptavidin-APC-Cy7. MuSC were isolated with BD Biosciences FACSAria II cell sorter as CD45^-^, CD11b^-^, CD31^-^, Sca1^-^, CD34^+^ and integrin α-7^+^ population. For the co-culture experiments, Fibro/adipogenic progenitors FAPs (Sca1^+^ and CD34^+^ population) were also recovered.

#### MuSC Culture

All cells were cultured in incubators at 37°C and 5% CO_2_. After isolation MuSC were plated on laminin (catalog number: 11243217001, Roche, 1:25 dilution in PBS) in growing medium (45% DMEM (catalog number: 10313-21, Gibco), 15% FBS (catalog number: FB-11, Omega Scientific), 1% Pen/Strep (catalog number: 15140163, Life technologies, Gibco^®^), 2.5 μg/mL FGFb (catalog number: 100-18B, Peprotech). For the knockdown experiments, cells were seeded at 3,000 cell/well and cultured for 72 hours before fixation, as described in the paragraph below. In each condition cells were further processed for immunostaining analysis.

#### Lentiviral infection and recombinant TnC treatment

Lentiviral particles were purchased from the MISSION shRNA library (Sigma). Tissue culture plates (96-well) were coated with laminin (20 mg/mL) for 1 hour at 37°C, followed by RetroNectin® (catalog number: T100A, Takara) (20 mg/mL in sterile PBS) for 2 hours at 37°C. Upon plating (cell density 3,000 cells/well), MuSC were treated with cationic polymer (hexadimethrine bromide, catalog number: 50-187-2421, Fisher) (8 µg/mL) at 37°C for 5 mins immediately followed by infection with 100 MOI of either custom control lentiviral GFP or lentiviral GFP sh-AnxA2 (trcn0000110696) in growth medium for 4 hours at 37°C and 5% CO_2_. At 4 hours, the media containing the lentiviral particles was removed and replace with growth medium. At 24 hours from the infection, half of the control and sh-AnxA2 samples were incubated with recombinant TnC (catalog number: 3358TC050, R&D) (5 µg/mL) or with the vehicle (PBS) for 48 hours in growth media at 37°C and 5% CO_2_. At the end of the experiment, samples were processed for: 1. mRNA extraction and qPCR to assess the knockdown (KD) efficiency; 2. Immunofluorescence. Only Pax7+ nuclei in GFP+ cells were counted and then normalized on the number of Pax7+ nuclei in control, non-treated cells.

#### Co-culture assay of MuSC and FAPs

After FACS isolation, cells were seeded on laminin coated 96-well plates at 2,000 cell/well per cell type (4,000 cell/well total) for co-culture. Control monocultures were seeded at 3,000 cell/well, as described above, and treated with recombinant TnC or vehicle (sterile PBS) to obtain a baseline for TnC effect on stem cell maintenance. Cells were cultured for 72 hours and then fixed with 4% PFA at room temperature for 10 mins. The cell samples were then either stored at 4°C or prepared for immunofluorescence. The percentage of Pax7+ cells in each co-culture combination was obtained by dividing the number of Pax7+ nuclei by the number of total nuclei minus the number of Pdgfra+ cells. All co-culture data was normalized on non-treated, monocultured WT MuSC. A minimum of 8 fields of view acquired with 20X objective were analyzed.

#### Skeletal muscle injury

Muscle injury was performed in tibialis anterior (TA). The protocol employed has been previously described in ^88^, with minor changes. Mice were anesthetized with isofluorane and ‘‘stabbed’’ 10 times before injecting 50 mL barium chloride (catalog number: 202738, Sigma) suspended in PBS (to 1.2% w/v) in several locations to distribute the solution to the entire tissue. MuSC and FAPs were isolated at 3 and 7 days after performing the injury for qPCR, or TA muscles were harvested at different timepoints and included in OCT (catalog number: 4583, VWR) for sectioning and immunohistochemistry.

#### *In vivo* EdU treatment

Mice were treated with EdU to assess for cell proliferation. With the use of a 29G needle insulin syringe, mice were intraperitoneally administered with 200μl of EdU (5mg/mL). Muscle groups of interested were then harvested 24 hours after injection.

#### Myofiber isolation for quiescent projection length quantification

Myofibers from EDL muscles of TnC-KO and WT adult mice were isolated following the protocol described by Kann *et al.*, 2022 ^52^, with minor adaptations. Mice were euthanized following the guidelines of approved AUF protocols. EDL muscles were harvested (by first severing the proximal tendon) and subjected to enzymatic dissociation (collagenase type II, catalog number: LS004177, Worthington) (700 units/mL) in low glucose (1 mg/mL) DMEM at 37°C for 55 mins. Dissociated single myofibers were manually collected and purified under a dissection microscope, then placed in the incubator to straighten for 10 mins. Myofibers were then fixed in 2% PFA at room temperature for 10 mins, subsequently washed twice in PBS, and stored in PBS at 4°C. All steps were performed in the presence of Y-27632 Rock inhibitor (catalog number: 72304, Stem Cell Technologies) (final concentration 10 µM) as described by Kann *et al*. 2022 ^52^.

#### Immunofluorescence

Muscle tissues (TA muscles) were isolated from healthy and injured mice at different time points. Tissues were embedded in OCT and frozen in 2-methyl butane. Tissues were sectioned in 10 μm thick slices and further processed through immunostaining. Fixation was performed with 4% PFA (catalog number: sc-281692, Santa Cruz Biotechnology). Tissue sections were washed in PBS twice, then permeabilized with 0.5% Triton 100-X (catalog number: 1003477329, EMD Millipore) in PBS and blocked in 10% goat serum (catalog number: 16210-072, Life technologies, Gibco^®^) and 0.1% Triton 100-X in PBS at room temperature for 1 hour. For Pax7 staining, samples were incubated with AffiniPure Fab fragment goat anti-mouse IgG (1:40, catalog number: 115-007-003, Jackson ImmunoResearch) solution in 0.2 µm filtered PBS at room temperature for 30 mins, then washed with PBS, and incubated in antigen retrieval (1:100, antigen unmasking solution, citric acid based, catalog number: H-3300, Vector Laboratories) solution at 92°C for 10 mins. Incubation with the primary antibodies was conducted at room temperature for 1 hour or at 4°C overnight in blocking buffer. All washes after incubation with antibodies were done by using PBS with 0.5% Triton 100-X. Myofibers were blocked in 10% goat serum with 0.3% Triton 100-X buffer at room temperature for 1 hour, followed by incubation with primary antibody blocking solution overnight at 4°C. Isolated cells were fixed in 4% PFA, washed in PBS twice, permeabilized at room temperature for 8 mins with 0.5% Triton 100-X, and incubated in 4% BSA (catalog number: SH30574.02, HyClone) with 0.5% Triton 100-X blocking buffer at room temperature for 1 hour or at 4°C overnight in blocking buffer. The primary antibodies used are the following: mouse anti-Pax7 (catalog number: Pax7-c, Developmental Studies Hybridoma Bank (DSHB), 1:50 dilution for cultured cells, 1:10 for tissue sections and myofibers), mouse anti-myogenin (catalog number: 556358, BD Biosciences; 1:100 dilution), mouse anti-MHC (catalog number: Mf20-c, DSHB,1:50 dilution), mouse anti-myosin (embryonic) (catalog number: F1.652, DSHB, 1:100 dilution), rabbit anti-tenascin-C (catalog number: AB19013, Millipore Sigma), rabbit anti-ɑ-tubulin (catalog number: ab18251, Abcam), rabbit anti-Pdgfra (catalog number: 3174T, Cell Signaling), rabbit anti-MyoD (catalog number: sc-760, Santa Cruz, 1:100 dilution), rabbit anti-laminin (catalog number: L9393, Sigma, 1:100 dilution), rat anti-laminin (catalog number: 05-206, Millipore, 1:100 dilution), chicken anti-GFP (catalog number: QAB10251, enquire Bio, 1:200 dilution). Alexa-conjugated secondary antibodies (Invitrogen, 1:500 dilution) were diluted in appropriate blocking buffer depending on the type of sample and incubated at room temperature for 45-60 mins. Nuclear DNA was stained with DAPI (Catalog number: MBD0011, Sigma). F-actin was labeled by using Alexa Fluor^TM^ 488 Phalloidin (catalog number: A12379, Life Technologies). Tissue section and myofibers were prepared for imaging in Fluoromount-G® (catalog number: 0100-01, SouthernBiotech) mounting solution. Images were acquired with Inverted IX81 Olympus Compound Fluorescence Microscope, XYZ Automated stage - ASI 2000 (Applied Scientific Instrumentation Inc.), with Color/monochrome cooled CCD camera - Spot RT3 and MetaMorph 7.11 Software (UIC, Molecular Devices) at 10X or 20X magnification or using confocal scanning through Leica TCS SP8 and LAS X software at 20X or 63X magnification. Leica DMi8 epifluorescent microscope was used for cell culture and tissue section imaging; a minimum of 8 fields of view acquired with 20X objective were analyzed. Nikon A1R HD confocal (running Nikon Elements software Version 5.42.04) with oil immersion 63X objective was used to acquire images of Pax7^+^ cells on myofibers. Sirius red staining visualization was obtained with Aperio AT2 Leica slide scanning system. All images were edited and modifications applied to the whole image through Fiji or Photoshop CS4 and Photoshop 2024 (Adobe).

#### Sirius Red staining of muscle sections

Tissue sections were re-hydrated for 2 mins in distilled water before nuclear staining with hematoxylin solution incubation for 5 mins. Sections were then washed for 10 mins with distilled water, and then incubated with Sirius red solution in picric acid for 1 hour. A 10 min wash was done with distilled water and subsequently with an acetic acid-based solution in distilled water for 2 mins. Three washes with 100% Ethanol were performed, each for 3 mins. The ethanol was removed before adding xylene for 3 mins. After removing the xylene sections were mounted with mounting solution. All steps were performed at room temperature.

#### Quantification of Muscle Tissue Cross-Sectional Area (CSA)

CSA quantification was performed in automated manner using and internally developed Macro through ImageJ64 ^89^ or Muscle Morphometry ImageJ plugin developed by Anthony Sinadinos (https://drive.google.com/drive/folders/0B_bBI7SbDQhCR1MxNEVXSlhiekE?resourcekey=0-8wdIKyTc0OKlB7WN67JqIw) by using the laminin fluorescent signal channel.

CSA and number of regenerating myofibers at 5 DPI in the experiments involving geriatric mice were calculated by using the embryonic myosin heavy-chain (eMyHC) fluorescent signal channel. The area of each myofiber and their number in each field of view were obtained by converting the images into binary, then followed by the command “Analyze particles” limited to the set threshold value.

#### Quantification of Tubulin Directionality

Confocal images of myofibers acquired with 20X objective were analyzed with Fiji by running the plugin ‘directionality’ on ROI (75x75 µm) of α-tubulin channels. The ‘Fourier Component’ method was used to ‘read’ the fluorescent signal, and the Gaussian curve fit. To obtain information on cytoskeleton organization in myofibers, dispersion and goodness values were collected; the dispersion is the value (in degrees) that represents tubulin alignment (smaller values mean more alignment); the goodness is the value (%) that represents how well the direction of the tubulin signal fits on a Gaussian curve (fit model used by the plugin).

#### RNA Isolation and Quantitative PCR

Total RNA was isolated with RNeasy Micro Kit (catalog number: 74004, Qiagen) following the manufacturer instruction. RNA quantification was performed with Qubit RNA HS Assay Kit (catalog number: Q32852, Invitrogen). The RNA samples for RNASeq analysis were aliquoted and further processed by SBPMDI Genomic Facility. The samples for qPCR analysis were further converted into cDNA with SuperScript^®^ VILO cDNA Synthesis Kit and Master Mix (catalog number: 11754050, Invitrogen) or High-Capacity cDNA Reverse Transcription Kit (catalog number: 4368814, Applied Biosystems) following manufacturer instructions. Real time PCR was performed on LightCycler^®^ 96 System (Roche) with Power SYBR**^®^** Green PCR Master Mix (catalog number: 4367659, Applied Biosystems), 5 µM primers concentration, and 0.5 ng of cDNA. Relative gene expression was calculated dividing the Ct value of each gene by the Ct value of the control (Large Ribosomal Protein, Rplp0, or Ribosomal Protein L7, Rpl7). The used primers are the followings:

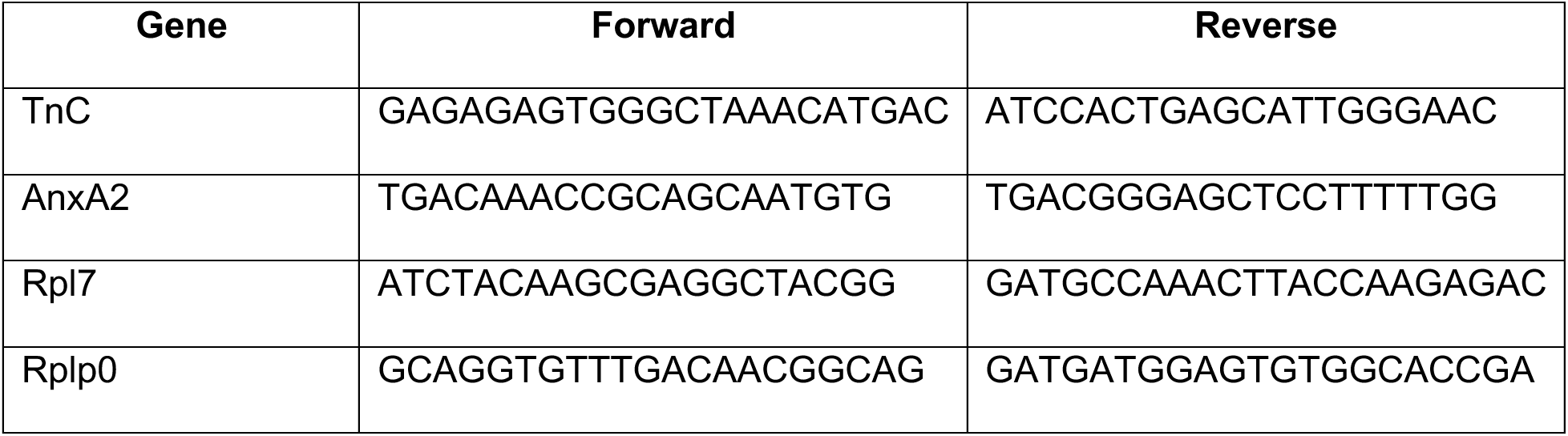

#### Co-immunoprecipitation (Co-IP)

To induce expression of TnC, mice were anesthetized with isofluorane and barium chloride (concentration described in the *skeletal muscle injury* section of the methods) was injected into the TA muscles. At 5 DPI, TA muscles were harvested, washed in 1X PBS for 5 min, snap frozen in liquid nitrogen, and immediately processed for protein lysate. Frozen TA muscles were lysed in lysis buffer (10X; 50 mM Tris, 100 mM NaCl, 1 mM EDTA, 0.5% NP40 + freshly added 1X phosphatases and proteases inhibitors, catalog numbers respectively: 04906837001 and 11836153001, Roche). Samples were kept at 4°C for 15 min for the lysis step, followed by centrifugation at 4°C for 15 min. The supernatant fraction was quantified by BSA standard curve before performing co-IP. Four mg of protein lysate were pre-cleared with protein G beads (catalog number: 10004D, Invitrogen) at 4°C for 1 hour, followed by incubation with anti-TnC antibody 1.5 µg/IP (catalog number: 10337, Immuno-Biological Laboratories) conjugated magnetic protein G beads at 4°C overnight on rotator. Prior to performing the Western blot assay, beads were collected using a magnetic rack and washed 3 times with 1X lysis buffer and 1 time with ammonium bicarbonate 50 mM at 4°C for 5 min on rotator. Beads were resuspended in 40 µL final volume of Western blot loading buffer, and Western blot was performed.

#### Western blot

Total protein lysates were prepared from cultures using lysis buffer (50 mM Tris HCl, 100 mM NaCl-, 1 mM EDTA, 1% Triton 100X, pH = 7.5) containing protease and phosphatase inhibitors. Cell membranes were removed by centrifugation and protein concentration was determined by using a BSA standard curve. Protein lysates or co-IP processed samples were loaded onto a NuPAGE 4-12% Bis-Tris gel and electrophoresis was performed in MOPS SDS running buffer. Proteins were transferred to a PVDF membrane and blocked with 5% BSA in PBST (PBS with 0.1% Triton 100X). Incubation with primary antibodies was performed at 4°C overnight. The antibodies used were: mouse anti-tenascin-(catalog number: 10337, Immuno-Biological Laboratories), rabbit anti-Annexin A2 (catalog number: 8235S, Cell Signaling Technology), mouse anti-vinculin (catalog number: sc-25336, Santa Cruz Biotechnology) and HRP-conjugated secondary antibodies (Santa Cruz), at the final concentration recommended by the manufacturers. Membranes were visualized with enhanced chemiluminescence (Pierce, Thermo Scientific Cat #32106) and developed on film.

#### Analysis of Single-Cell RNA sequencing

scRNA-seq data was downloaded from Oprescu *et al.*, 2020 ^54^ using the accession number GSE138826. Bioinformatics analysis was performed using Seurat (version 5.0.2 and R version 4.3.3), dplyr, reshape2, tidyverse, and scCustomize. Visualizations were performed using the Seurat function (VlnPlot), the CellChat package (see Cell-Cell Communication Analysis below), and the R packages ggplot2 and dittoSeq.

#### Cell-Cell Communication Analysis

The cell-cell communication and visualization were performed using CellChat (version 2.1.0). CellChatDB is a manually curated database of literature-supported ligand-receptor interactions in multiple species (we sub-selected the CellChatDB.mouse database), including multi-subunit structure of ligand-receptor complexes and co-factors. We manually updated the CellChatDB.mouse interaction database to include several known interactions that were not included by default (TNC_ANXA2, TNC_ITGA7_ITGB1, TNC_EGFR, TNC_ITGAV_ITGB1, TNC_ITGA2_ITGB1, and TNC_TLR4) after conducting a thorough literature search. For the cell– interaction analysis, the expression levels were calculated relative to the total read mapping to the same set of coding genes in all transcriptomes. The expression values were averaged within each single-cell cluster/cell sample. Visualizations were performed using the following CellChat functions: netVisual_aggregate, netVisual_heatmap, netVisual_bubble, and netAnalysis_individual.

#### Statistical Analysis

Error bars in the figures represent standard error of the mean and number of biological replicates is indicated by n, while number of cells or fields of view are indicated respectively with N_cell_ or N_fov_ in the figure legends. Statistical significance was tested with Student’s *t* test for two groups comparison, and either one-way ANOVA or two-way ANOVA for multiple groups. P values <0.05 were considered significant. All the statistical analysis was performed with Prism 7 program (GraphPad).

## Notes

### Competing Interest Statement

The authors have declared no competing interest.

## REFERENCES

1. Dumont, N.A., Wang, Y.X., and Rudnicki, M.A. (2015). Intrinsic and extrinsic mechanisms regulating satellite cell function. Development 142, 1572–1581. 10.1242/dev.114223.

2. Yin, Y., He, G.J., Hu, S., Tse, E.H.Y., and Cheung, T.H. (2024). Muscle stem cell niche dynamics during muscle homeostasis and regeneration. Curr Top Dev Biol 158, 151–177. 10.1016/bs.ctdb.2024.02.008.

3. Majchrzak, K., Hentschel, E., Honzke, K., Geithe, C., and von Maltzahn, J. (2024). We need to talk-how muscle stem cells communicate. Front Cell Dev Biol 12, 1378548. 10.3389/fcell.2024.1378548.

4. Joe, A.W., Yi, L., Natarajan, A., Le Grand, F., So, L., Wang, J., Rudnicki, M.A., and Rossi, F.M. (2010). Muscle injury activates resident fibro/adipogenic progenitors that facilitate myogenesis. Nat Cell Biol 12, 153–163. 10.1038/ncb2015.

5. Uezumi, A., Fukada, S., Yamamoto, N., Takeda, S., and Tsuchida, K. (2010). Mesenchymal progenitors distinct from satellite cells contribute to ectopic fat cell formation in skeletal muscle. Nat Cell Biol 12, 143–152. 10.1038/ncb2014.

6. Wosczyna, M.N., Konishi, C.T., Perez Carbajal, E.E., Wang, T.T., Walsh, R.A., Gan, Q., Wagner, M.W., and Rando, T.A. (2019). Mesenchymal Stromal Cells Are Required for Regeneration and Homeostatic Maintenance of Skeletal Muscle. Cell Rep 27, 2029–2035 e2025. 10.1016/j.celrep.2019.04.074.

7. Malecova, B., Gatto, S., Etxaniz, U., Passafaro, M., Cortez, A., Nicoletti, C., Giordani, L., Torcinaro, A., De Bardi, M., Bicciato, S., et al. (2018). Dynamics of cellular states of fibro-adipogenic progenitors during myogenesis and muscular dystrophy. Nat Commun 9, 3670. 10.1038/s41467-018-06068-6.

8. Contreras, O., Cruz-Soca, M., Theret, M., Soliman, H., Tung, L.W., Groppa, E., Rossi, F.M., and Brandan, E. (2019). Cross-talk between TGF-beta and PDGFRalpha signaling pathways regulates the fate of stromal fibro-adipogenic progenitors. J Cell Sci 132. 10.1242/jcs.232157.

9. Uapinyoying, P., Hogarth, M., Battacharya, S., Mazala, D.A.G., Panchapakesan, K., Bonnemann, C.G., and Jaiswal, J.K. (2023). Single-cell transcriptomic analysis of the identity and function of fibro/adipogenic progenitors in healthy and dystrophic muscle. iScience 26, 107479. 10.1016/j.isci.2023.107479.

10. Kotsaris, G., Qazi, T.H., Bucher, C.H., Zahid, H., Pohle-Kronawitter, S., Ugorets, V., Jarassier, W., Borno, S., Timmermann, B., Giesecke-Thiel, C., et al. (2023). Odd skipped-related 1 controls the pro-regenerative response of fibro-adipogenic progenitors. NPJ Regen Med 8, 19. 10.1038/s41536-023-00291-6.

11. Wang, K., Yang, J., An, Y., Wang, J., Tan, S., Xu, H., and Dong, Y. (2024). MST1/2 regulates fibro/adipogenic progenitor fate decisions in skeletal muscle regeneration. Stem Cell Reports 19, 501–514. 10.1016/j.stemcr.2024.02.010.

12. Molina, T., Fabre, P., and Dumont, N.A. (2021). Fibro-adipogenic progenitors in skeletal muscle homeostasis, regeneration and diseases. Open Biol 11, 210110. 10.1098/rsob.210110.

13. Helzer, D., Kannan, P., Reynolds, J.C., Gibbs, D.E., and Crosbie, R.H. (2024). Role of microenvironment on muscle stem cell function in health, adaptation, and disease. Curr Top Dev Biol 158, 179–201. 10.1016/bs.ctdb.2024.02.002.

14. Loreti, M., and Sacco, A. (2022). The jam session between muscle stem cells and the extracellular matrix in the tissue microenvironment. NPJ Regen Med 7, 16. 10.1038/s41536-022-00204-z.

15. Settles, D.L., Cihak, R.A., and Erickson, H.P. (1996). Tenascin-C expression in dystrophin-related muscular dystrophy. Muscle Nerve 19, 147–154. 10.1002/(SICI)1097-4598(199602)19:2<147::AID-MUS4>3.0.CO;2-E.

16. Gullberg, D., Velling, T., Sjoberg, G., Salmivirta, K., Gaggero, B., Tiger, C.F., Edstrom, L., and Sejersen, T. (1997). Tenascin-C expression correlates with macrophage invasion in Duchenne muscular dystrophy and in myositis. Neuromuscul Disord 7, 39–54. 10.1016/s0960-8966(96)00391-4.

17. Fluck, M., Chiquet, M., Schmutz, S., Mayet-Sornay, M.H., and Desplanches, D. (2003). Reloading of atrophied rat soleus muscle induces tenascin-C expression around damaged muscle fibers. Am J Physiol Regul Integr Comp Physiol 284, R792–801. 10.1152/ajpregu.00060.2002.

18. Tierney, M.T., Gromova, A., Sesillo, F.B., Sala, D., Spenle, C., Orend, G., and Sacco, A. (2016). Autonomous Extracellular Matrix Remodeling Controls a Progressive Adaptation in Muscle Stem Cell Regenerative Capacity during Development. Cell Rep 14, 1940–1952. 10.1016/j.celrep.2016.01.072.

19. Kardon, G. (1998). Muscle and tendon morphogenesis in the avian hind limb. Development 125, 4019–4032. 10.1242/dev.125.20.4019.

20. Zhou, S., Zhang, W., Cai, G., Ding, Y., Wei, C., Li, S., Yang, Y., Qin, J., Liu, D., Zhang, H., et al. (2020). Myofiber necroptosis promotes muscle stem cell proliferation via releasing Tenascin-C during regeneration. Cell Res 30, 1063–1077. 10.1038/s41422-020-00393-6.

21. Proietti, D., Giordani, L., De Bardi, M., D’Ercole, C., Lozanoska-Ochser, B., Amadio, S., Volonte, C., Marinelli, S., Muchir, A., Bouche, M., et al. (2021). Activation of skeletal muscle-resident glial cells upon nerve injury. JCI Insight 6. 10.1172/jci.insight.143469.

22. Nicoletti, C., Wei, X., Etxaniz, U., D’Ercole, C., Madaro, L., Perera, R., and Puri, P.L. (2023). Muscle denervation promotes functional interactions between glial and mesenchymal cells through NGFR and NGF. iScience 26, 107114. 10.1016/j.isci.2023.107114.

23. Lukjanenko, L., Jung, M.J., Hegde, N., Perruisseau-Carrier, C., Migliavacca, E., Rozo, M., Karaz, S., Jacot, G., Schmidt, M., Li, L., et al. (2016). Loss of fibronectin from the aged stem cell niche affects the regenerative capacity of skeletal muscle in mice. Nat Med 22, 897–905. 10.1038/nm.4126.

24. Nakao, N., Hiraiwa, N., Yoshiki, A., Ike, F., and Kusakabe, M. (1998). Tenascin-C promotes healing of Habu-snake venom-induced glomerulonephritis: studies in knockout congenic mice and in culture. Am J Pathol 152, 1237–1245.

25. Koyama, Y., Kusubata, M., Yoshiki, A., Hiraiwa, N., Ohashi, T., Irie, S., and Kusakabe, M. (1998). Effect of tenascin-C deficiency on chemically induced dermatitis in the mouse. J Invest Dermatol 111, 930–935. 10.1046/j.1523-1747.1998.00401.x.

26. Kiernan, B.W., Garcion, E., Ferguson, J., Frost, E.E., Torres, E.M., Dunnett, S.B., Saga, Y., Aizawa, S., Faissner, A., Kaur, R., et al. (1999). Myelination and behaviour of tenascin-C null transgenic mice. Eur J Neurosci 11, 3082–3092. 10.1046/j.1460-9568.1999.00729.x.

27. Tamaoki, M., Imanaka-Yoshida, K., Yokoyama, K., Nishioka, T., Inada, H., Hiroe, M., Sakakura, T., and Yoshida, T. (2005). Tenascin-C regulates recruitment of myofibroblasts during tissue repair after myocardial injury. Am J Pathol 167, 71–80. 10.1016/S0002-9440(10)62954-9.

28. de Chevigny, A., Lemasson, M., Saghatelyan, A., Sibbe, M., Schachner, M., and Lledo, P.M. (2006). Delayed onset of odor detection in neonatal mice lacking tenascin-C. Mol Cell Neurosci 32, 174–186. 10.1016/j.mcn.2006.04.002.

29. Okamura, N., Hasegawa, M., Nakoshi, Y., Iino, T., Sudo, A., Imanaka-Yoshida, K., Yoshida, T., and Uchida, A. (2010). Deficiency of tenascin-C delays articular cartilage repair in mice. Osteoarthritis Cartilage 18, 839–848. 10.1016/j.joca.2009.08.013.

30. Sumioka, T., Fujita, N., Kitano, A., Okada, Y., and Saika, S. (2011). Impaired angiogenic response in the cornea of mice lacking tenascin C. Invest Ophthalmol Vis Sci 52, 2462–2467. 10.1167/iovs.10-5750.

31. Ohta, M., Sakai, T., Saga, Y., Aizawa, S., and Saito, M. (1998). Suppression of hematopoietic activity in tenascin-C-deficient mice. Blood 91, 4074–4083.

32. Nakamura-Ishizu, A., Okuno, Y., Omatsu, Y., Okabe, K., Morimoto, J., Uede, T., Nagasawa, T., Suda, T., and Kubota, Y. (2012). Extracellular matrix protein tenascin-C is required in the bone marrow microenvironment primed for hematopoietic regeneration. Blood 119, 5429–5437. 10.1182/blood-2011-11-393645.

33. Fluck, M., Mund, S.I., Schittny, J.C., Klossner, S., Durieux, A.C., and Giraud, M.N. (2008). Mechano-regulated tenascin-C orchestrates muscle repair. Proc Natl Acad Sci U S A 105, 13662–13667. 10.1073/pnas.0805365105.

34. Midwood, K.S., Hussenet, T., Langlois, B., and Orend, G. (2011). Advances in tenascin-C biology. Cell Mol Life Sci 68, 3175–3199. 10.1007/s00018-011-0783-6.

35. Midwood, K.S., Chiquet, M., Tucker, R.P., and Orend, G. (2016). Tenascin-C at a glance. J Cell Sci 129, 4321–4327. 10.1242/jcs.190546.

36. Chung, C.Y., and Erickson, H.P. (1994). Cell surface annexin II is a high affinity receptor for the alternatively spliced segment of tenascin-C. J Cell Biol 126, 539–548. 10.1083/jcb.126.2.539.

37. Wang, Z., Wei, Q., Han, L., Cao, K., Lan, T., Xu, Z., Wang, Y., Gao, Y., Xue, J., Shan, F., et al. (2018). Tenascin-c renders a proangiogenic phenotype in macrophage via annexin II. J Cell Mol Med 22, 429–438. 10.1111/jcmm.13332.

38. Chung, C.Y., Murphy-Ullrich, J.E., and Erickson, H.P. (1996). Mitogenesis, cell migration, and loss of focal adhesions induced by tenascin-C interacting with its cell surface receptor, annexin II. Mol Biol Cell 7, 883–892. 10.1091/mbc.7.6.883.

39. Demonbreun, A.R., Quattrocelli, M., Barefield, D.Y., Allen, M.V., Swanson, K.E., and McNally, E.M. (2016). An actin-dependent annexin complex mediates plasma membrane repair in muscle. J Cell Biol 213, 705–718. 10.1083/jcb.201512022.

40. Hogarth, M.W., Defour, A., Lazarski, C., Gallardo, E., Diaz Manera, J., Partridge, T.A., Nagaraju, K., and Jaiswal, J.K. (2019). Fibroadipogenic progenitors are responsible for muscle loss in limb girdle muscular dystrophy 2B. Nat Commun 10, 2430. 10.1038/s41467-019-10438-z.

41. Kayejo, V.G., Fellner, H., Thapa, R., and Keyel, P.A. (2023). Translational implications of targeting annexin A2: From membrane repair to muscular dystrophy, cardiovascular disease and cancer. Clin Transl Discov 3. 10.1002/ctd2.240.

42. Fuchs, E., and Blau, H.M. (2020). Tissue Stem Cells: Architects of Their Niches. Cell Stem Cell 27, 532–556. 10.1016/j.stem.2020.09.011.

43. Li, E.W., McKee-Muir, O.C., and Gilbert, P.M. (2018). Cellular Biomechanics in Skeletal Muscle Regeneration. Curr Top Dev Biol 126, 125–176. 10.1016/bs.ctdb.2017.08.007.

44. Mashinchian, O., Pisconti, A., Le Moal, E., and Bentzinger, C.F. (2018). The Muscle Stem Cell Niche in Health and Disease. Curr Top Dev Biol 126, 23–65. 10.1016/bs.ctdb.2017.08.003.

45. Lukjanenko, L., Karaz, S., Stuelsatz, P., Gurriaran-Rodriguez, U., Michaud, J., Dammone, G., Sizzano, F., Mashinchian, O., Ancel, S., Migliavacca, E., et al. (2019). Aging Disrupts Muscle Stem Cell Function by Impairing Matricellular WISP1 Secretion from Fibro-Adipogenic Progenitors. Cell Stem Cell 24, 433–446 e437. 10.1016/j.stem.2018.12.014.

46. Uezumi, A., Ikemoto-Uezumi, M., Zhou, H., Kurosawa, T., Yoshimoto, Y., Nakatani, M., Hitachi, K., Yamaguchi, H., Wakatsuki, S., Araki, T., et al. (2021). Mesenchymal Bmp3b expression maintains skeletal muscle integrity and decreases in age-related sarcopenia. J Clin Invest 131. 10.1172/JCI139617.

47. Forsberg, E., Hirsch, E., Frohlich, L., Meyer, M., Ekblom, P., Aszodi, A., Werner, S., and Fassler, R. (1996). Skin wounds and severed nerves heal normally in mice lacking tenascin-C. Proc Natl Acad Sci U S A 93, 6594–6599. 10.1073/pnas.93.13.6594.

48. Kassar-Duchossoy, L., Giacone, E., Gayraud-Morel, B., Jory, A., Gomes, D., and Tajbakhsh, S. (2005). Pax3/Pax7 mark a novel population of primitive myogenic cells during development. Genes Dev 19, 1426–1431. 10.1101/gad.345505.

49. Relaix, F., Montarras, D., Zaffran, S., Gayraud-Morel, B., Rocancourt, D., Tajbakhsh, S., Mansouri, A., Cumano, A., and Buckingham, M. (2006). Pax3 and Pax7 have distinct and overlapping functions in adult muscle progenitor cells. J Cell Biol 172, 91–102. 10.1083/jcb.200508044.

50. Vasyutina, E., Lenhard, D.C., Wende, H., Erdmann, B., Epstein, J.A., and Birchmeier, C. (2007). RBP-J (Rbpsuh) is essential to maintain muscle progenitor cells and to generate satellite cells. Proc Natl Acad Sci U S A 104, 4443–4448. 10.1073/pnas.0610647104.

51. Chakkalakal, J.V., Christensen, J., Xiang, W., Tierney, M.T., Boscolo, F.S., Sacco, A., and Brack, A.S. (2014). Early forming label-retaining muscle stem cells require p27kip1 for maintenance of the primitive state. Development 141, 1649–1659. 10.1242/dev.100842.

52. Kann, A.P., Hung, M., Wang, W., Nguyen, J., Gilbert, P.M., Wu, Z., and Krauss, R.S. (2022). An injury-responsive Rac-to-Rho GTPase switch drives activation of muscle stem cells through rapid cytoskeletal remodeling. Cell Stem Cell 29, 933–947 e936. 10.1016/j.stem.2022.04.016.

53. Ma, N., Chen, D., Lee, J.H., Kuri, P., Hernandez, E.B., Kocan, J., Mahmood, H., Tichy, E.D., Rompolas, P., and Mourkioti, F. (2022). Piezo1 regulates the regenerative capacity of skeletal muscles via orchestration of stem cell morphological states. Sci Adv 8, eabn0485. 10.1126/sciadv.abn0485.

54. Oprescu, S.N., Yue, F., Qiu, J., Brito, L.F., and Kuang, S. (2020). Temporal Dynamics and Heterogeneity of Cell Populations during Skeletal Muscle Regeneration. iScience 23, 100993. 10.1016/j.isci.2020.100993.

55. Jin, S., Guerrero-Juarez, C.F., Zhang, L., Chang, I., Ramos, R., Kuan, C.H., Myung, P., Plikus, M.V., and Nie, Q. (2021). Inference and analysis of cell-cell communication using CellChat. Nat Commun 12, 1088. 10.1038/s41467-021-21246-9.

56. Jin S, P.M., Nie Q (2023). CellChat for systematic analysis of cell-cell communication from single-cell and spatially resolved transcriptomics. In I. University of California, ed.

57. Chiquet-Ehrismann, R., and Tucker, R.P. (2011). Tenascins and the importance of adhesion modulation. Cold Spring Harb Perspect Biol 3. 10.1101/cshperspect.a004960.

58. Chiquet-Ehrismann, R., Orend, G., Chiquet, M., Tucker, R.P., and Midwood, K.S. (2014). Tenascins in stem cell niches. Matrix Biol 37, 112–123. 10.1016/j.matbio.2014.01.007.

59. Abedsaeidi, M., Hojjati, F., Tavassoli, A., and Sahebkar, A. (2024). Biology of Tenascin C and its Role in Physiology and Pathology. Curr Med Chem 31, 2706–2731. 10.2174/0929867330666230404124229.

60. Brack, A.S., Bildsoe, H., and Hughes, S.M. (2005). Evidence that satellite cell decrement contributes to preferential decline in nuclear number from large fibres during murine age-related muscle atrophy. J Cell Sci 118, 4813–4821. 10.1242/jcs.02602.

61. Shefer, G., Van de Mark, D.P., Richardson, J.B., and Yablonka-Reuveni, Z. (2006). Satellite-cell pool size does matter: defining the myogenic potency of aging skeletal muscle. Dev Biol 294, 50–66. 10.1016/j.ydbio.2006.02.022.

62. Collins, C.A., Zammit, P.S., Ruiz, A.P., Morgan, J.E., and Partridge, T.A. (2007). A population of myogenic stem cells that survives skeletal muscle aging. Stem Cells 25, 885–894. 10.1634/stemcells.2006-0372.

63. Webster, M.T., Manor, U., Lippincott-Schwartz, J., and Fan, C.M. (2016). Intravital Imaging Reveals Ghost Fibers as Architectural Units Guiding Myogenic Progenitors during Regeneration. Cell Stem Cell 18, 243–252. 10.1016/j.stem.2015.11.005.

64. Verma, M., Asakura, Y., Murakonda, B.S.R., Pengo, T., Latroche, C., Chazaud, B., McLoon, L.K., and Asakura, A. (2018). Muscle Satellite Cell Cross-Talk with a Vascular Niche Maintains Quiescence via VEGF and Notch Signaling. Cell Stem Cell 23, 530–543 e539. 10.1016/j.stem.2018.09.007.

65. Haroon, M., Klein-Nulend, J., Bakker, A.D., Jin, J., Seddiqi, H., Offringa, C., de Wit, G.M.J., Le Grand, F., Giordani, L., Liu, K.J., et al. (2021). Myofiber stretch induces tensile and shear deformation of muscle stem cells in their native niche. Biophys J 120, 2665–2678. 10.1016/j.bpj.2021.05.021.

66. Murphy-Ullrich, J.E., Lightner, V.A., Aukhil, I., Yan, Y.Z., Erickson, H.P., and Hook, M. (1991). Focal adhesion integrity is downregulated by the alternatively spliced domain of human tenascin. J Cell Biol 115, 1127–1136. 10.1083/jcb.115.4.1127.

67. Midwood, K.S., and Schwarzbauer, J.E. (2002). Tenascin-C modulates matrix contraction via focal adhesion kinase- and Rho-mediated signaling pathways. Mol Biol Cell 13, 3601–3613. 10.1091/mbc.e02-05-0292.

68. Sun, Z., Schwenzer, A., Rupp, T., Murdamoothoo, D., Vegliante, R., Lefebvre, O., Klein, A., Hussenet, T., and Orend, G. (2018). Tenascin-C Promotes Tumor Cell Migration and Metastasis through Integrin alpha9beta1-Mediated YAP Inhibition. Cancer Res 78, 950–961. 10.1158/0008-5472.CAN-17-1597.

69. Demonbreun, A.R., Fallon, K.S., Oosterbaan, C.C., Bogdanovic, E., Warner, J.L., Sell, J.J., Page, P.G., Quattrocelli, M., Barefield, D.Y., and McNally, E.M. (2019). Recombinant annexin A6 promotes membrane repair and protects against muscle injury. J Clin Invest 129, 4657–4670. 10.1172/JCI128840.

70. Gerke, V., Gavins, F.N.E., Geisow, M., Grewal, T., Jaiswal, J.K., Nylandsted, J., and Rescher, U. (2024). Annexins-a family of proteins with distinctive tastes for cell signaling and membrane dynamics. Nat Commun 15, 1574. 10.1038/s41467-024-45954-0.

71. Foltz, S.J., Cui, Y.Y., Choo, H.J., and Hartzell, H.C. (2021). ANO5 ensures trafficking of annexins in wounded myofibers. J Cell Biol 220. 10.1083/jcb.202007059.

72. Lennon, N.J., Kho, A., Bacskai, B.J., Perlmutter, S.L., Hyman, B.T., and Brown, R.H., Jr. (2003). Dysferlin interacts with annexins A1 and A2 and mediates sarcolemmal wound-healing. J Biol Chem 278, 50466–50473. 10.1074/jbc.M307247200.

73. Defour, A., Medikayala, S., Van der Meulen, J.H., Hogarth, M.W., Holdreith, N., Malatras, A., Duddy, W., Boehler, J., Nagaraju, K., and Jaiswal, J.K. (2017). Annexin A2 links poor myofiber repair with inflammation and adipogenic replacement of the injured muscle. Hum Mol Genet 26, 1979–1991. 10.1093/hmg/ddx065.

74. Bittel, D.C., Chandra, G., Tirunagri, L.M.S., Deora, A.B., Medikayala, S., Scheffer, L., Defour, A., and Jaiswal, J.K. (2020). Annexin A2 Mediates Dysferlin Accumulation and Muscle Cell Membrane Repair. Cells 9. 10.3390/cells9091919.

75. Babbin, B.A., Parkos, C.A., Mandell, K.J., Winfree, L.M., Laur, O., Ivanov, A.I., and Nusrat, A. (2007). Annexin 2 regulates intestinal epithelial cell spreading and wound closure through Rho-related signaling. Am J Pathol 170, 951–966. 10.2353/ajpath.2007.060647.

76. Matsuda, A., Tagawa, Y., Yamamoto, K., Matsuda, H., and Kusakabe, M. (1999). Identification and immunohistochemical localization of annexin II in rat cornea. Curr Eye Res 19, 368–375. 10.1076/ceyr.19.4.368.5306.

77. Rescher, U., Ludwig, C., Konietzko, V., Kharitonenkov, A., and Gerke, V. (2008). Tyrosine phosphorylation of annexin A2 regulates Rho-mediated actin rearrangement and cell adhesion. J Cell Sci 121, 2177–2185. 10.1242/jcs.028415.

78. Chiquet, M., and Fambrough, D.M. (1984). Chick myotendinous antigen. II. A novel extracellular glycoprotein complex consisting of large disulfide-linked subunits. J Cell Biol 98, 1937–1946. 10.1083/jcb.98.6.1937.

79. Mozzetta, C., Consalvi, S., Saccone, V., Tierney, M., Diamantini, A., Mitchell, K.J., Marazzi, G., Borsellino, G., Battistini, L., Sassoon, D., et al. (2013). Fibroadipogenic progenitors mediate the ability of HDAC inhibitors to promote regeneration in dystrophic muscles of young, but not old Mdx mice. EMBO Mol Med 5, 626–639. 10.1002/emmm.201202096.

80. Madaro, L., Passafaro, M., Sala, D., Etxaniz, U., Lugarini, F., Proietti, D., Alfonsi, M.V., Nicoletti, C., Gatto, S., De Bardi, M., et al. (2018). Denervation-activated STAT3-IL-6 signalling in fibro-adipogenic progenitors promotes myofibres atrophy and fibrosis. Nat Cell Biol 20, 917–927. 10.1038/s41556-018-0151-y.

81. Jinnin, M., Ihn, H., Asano, Y., Yamane, K., Trojanowska, M., and Tamaki, K. (2004). Tenascin-C upregulation by transforming growth factor-beta in human dermal fibroblasts involves Smad3, Sp1, and Ets1. Oncogene 23, 1656–1667. 10.1038/sj.onc.1207064.

82. Sarasa-Renedo, A., Tunc-Civelek, V., and Chiquet, M. (2006). Role of RhoA/ROCK-dependent actin contractility in the induction of tenascin-C by cyclic tensile strain. Exp Cell Res 312, 1361–1370. 10.1016/j.yexcr.2005.12.025.

83. Giblin, S.P., and Midwood, K.S. (2015). Tenascin-C: Form versus function. Cell Adh Migr 9, 48–82. 10.4161/19336918.2014.987587.

84. Garcia-Carrizo F, G.S., Lenihan-Geels G, Jank AM, Leer M, Soultoukis GA, Oveisi M, Herpich C, Garrido CA, Kotsaris G, Pohle-Kronawitter S, Tsamo-Tetou A, Graja A, Ost M, Villacorta L, Knecht RS, Klaus S, Schurmann A, Stricker S, Schmidt-Bleek K, Cipitria A, Duda GN, Benes V, Muller-Werdan U, Norman K, Schulz TJ (2023). Aging impairs skeletal muscle regeneration by promoting fibro/fatty degeneration and inhibiting inflammation resolution via fibro-adipogenic progenitors. In N. German Institute of Human Nutrition Potsdam-Rehbrücke, Germany, ed.

85. Albacete-Albacete, L., Navarro-Lerida, I., Lopez, J.A., Martin-Padura, I., Astudillo, A.M., Ferrarini, A., Van-Der-Heyden, M., Balsinde, J., Orend, G., Vazquez, J., and Del Pozo, M.A. (2020). ECM deposition is driven by caveolin-1-dependent regulation of exosomal biogenesis and cargo sorting. J Cell Biol 219. 10.1083/jcb.202006178.

86. Campos, A., Salomon, C., Bustos, R., Diaz, J., Martinez, S., Silva, V., Reyes, C., Diaz-Valdivia, N., Varas-Godoy, M., Lobos-Gonzalez, L., and Quest, A.F. (2018). Caveolin-1-containing extracellular vesicles transport adhesion proteins and promote malignancy in breast cancer cell lines. Nanomedicine (Lond) 13, 2597–2609. 10.2217/nnm-2018-0094.

87. Gromova, A., Tierney, M.T., and Sacco, A. (2015). FACS-based Satellite Cell Isolation From Mouse Hind Limb Muscles. Bio Protoc 5. 10.21769/bioprotoc.1558.

88. Tierney, M.T., and Sacco, A. (2016). Inducing and Evaluating Skeletal Muscle Injury by Notexin and Barium Chloride. Methods Mol Biol 1460, 53–60. 10.1007/978-1-4939-3810-0_5.

89. Schneider, C.A., Rasband, W.S., and Eliceiri, K.W. (2012). NIH Image to ImageJ: 25 years of image analysis. Nat Methods 9, 671–675. 10.1038/nmeth.2089.

