## Supplementary Figures and Legends for "Tenascin-C from the tissue microenvironment promotes muscle stem cell self-renewal through Annexin A2"

[1]Development, Aging and Regeneration Program, Sanford Burnham Prebys Medical Discovery Institute, 10901 N Torrey Pines Rd, La Jolla, CA 92037, USA
[2#]Current affiliation: J&J, 3880 Murphy Canyon Rd, San Diego, CA 92123, USA
[3#]Current affiliation: University of California San Diego, 9500 Gilman Drive, La Jolla, CA 92093, USA
[4#]Current affiliation: Avidity Biosciences, Inc., 10578 Science Center Drive Suite 125, San Diego, CA 92121, USA
[5]Rice University, 6100 Main St, Huston, TX 77005, USA

*These authors equally contributed to the manuscript

Corresponding author: Alessandra Sacco, Ph.D.
Development, Aging and Regeneration Program
Sanford Burnham Prebys Medical Discovery Institute
10901 North Torrey Pines Road
La Jolla, CA 92037, USA


#### SUPPLEMENTARY FIGURES

**Figure S1. TnC-KO mice do not exhibit a skeletal muscle phenotype at prenatal stages, related to Figure 1.**

(**A**) Comparison of $Pax7^+$ cell numbers (per $mm^2$) between postnatal stage P14 and P30 in WT and TnC-KO mice (n=4).

(**B**) Comparison of muscle growth (CSA, $\mu m^2$) between P14 and P30 in WT and TnC-KO mice (n≥4).

(**C**) Representative immunofluorescence (IF) images of hindlimb cross-sections from WT and TnC-KO mice (Pax7, green; Ki-67, red; laminin, gray; DAPI, blue) (scale bar= 50 μm) and quantification of $Pax7^+$ cell proliferation (Ki-67) during embryogenesis (E 16.5) (n=3).

(**D**) Representative immunofluorescence (IF) images of TA muscle cross-sections from WT and TnC-KO mice (Pax7, green; Ki-67, red; laminin, gray; DAPI, blue) (scale bar= 50 μm) and quantification of $Pax7^+$ cell proliferation (Ki-67) during postnatal growth (P14) (n≥3).

(**E**) Representative immunofluorescence (IF) images of homeostatic TA muscle cross-sections from adult (3-6 months old) WT and TnC-KO mice (Pax7, green; Ki-67, red; DAPI, blue) (scale bar= 50 μm) and quantification of $Pax7^+$ cell proliferation (Ki-67) (n=3).

(**F**) Representative immunofluorescence (IF) images of injured (5 DPI) TA muscle cross-sections from adult (3-6 months old) WT and TnC-KO mice (Pax7, green; EdU, red; laminin, gray; DAPI, blue) (scale bar= 50 μm) and quantification of $Pax7^+$ cell proliferation (EdU – timepoint 24 hours) (n=3).

Data are represented as mean ± SEM; $p>0.05$, $^{**}p<0.01$, $t$ test (**A, C, D, F**), $^{***}p<0.001$, one-way ANOVA (**B**).

**Figure S2. Tenocytes express TnC during homeostasis and regeneration, related to Figure 3.**

(**A**) TnC mRNA dynamics across regeneration in tenocytes (online available dataset).

(**B**) Tenascin signaling network heat map plot at 21 DPI (online available dataset).

(**C**) Chord diagrams representing incoming signaling to MuSC in uninjured and 21 DPI TA muscles (online available dataset).

**Figure S3. TnC is expressed by MuSC and FAPs at different levels depending on the presence of AnxA2 transcript, related to Figure 4.**

(**A**) Violin plots depicting the different expression dynamics of TnC at different DPIs (0, 3.5, 5, 21) in MuSC and FAPs (online available dataset).

(**B**) Table of the raw data of cell numbers per condition related to the dot plot in main Figure 4D representing the changes in the relative expression of TnC and the percentages of TnC expressing cell populations within $Pax7^+/AnxA2^+$ and $Pax7^+/AnxA2^-$ MuSC and $Pdgfra^+/AnxA2^+$ and $Pdgfra^+/AnxA2^-$ FAPs in uninjured TA muscles and across different timepoints (3.5, 5, 21 DPI).

(**C**) Dot plot representing the changes in the relative expression of TnC and the percentages of TnC expressing cell populations within $MyoD^+/AnxA2^+$ and $MyoD^+/AnxA2^-$ MuSC in uninjured TA muscles and across different timepoints (3.5, 5, 21 DPI), and corresponding table of the raw data of cell numbers per condition.

(**D**) Quantification of the efficiency of the AnxA2 KD via qPCR (n=3).

(**E**) Representative immunofluorescence images of GFP-control and AnxA2 KD being treated or not with the recombinant TnC (48hrs treatment) (GFP, green; MyoD, red; DAPI, blue) (scale bar = 50 μm) and quantification of MyoD+ cells normalized on non-treated GFP-control infected MuSC (n=3, $N_{fov}$= 30).

Data are represented as mean ± SEM; ****$p$<0.0001, $t$ test (**D**); $p$>0.05, one-way ANOVA (**E**).

**Figure S4. TnC-KO myofibers show a disorganized $\alpha$-tubulin network, related to Figure 5.**

(**A**) Representative immunofluorescence images of myofiber cytoskeleton organization in adult (4-6 months old) mice ($\alpha$-tubulin, magenta). Scale bar= 50 µm.

(**B**) Representative images of the output graphs obtained by using the Fiji Fourier transform 'directionality' tool.

(**C**) Comparison of the goodness of the fit, the Gaussian curve is the fit model implemented in this analysis, between WT and TnC-KO myofibers after running the 'directionality' tool. Dispersion (˚) values indicating the level of alignment of cytoskeletal $\alpha$-tubulin in WT and TnC-KO myofibers (n=3, $N_{cell}$= 75).

Data are represented as the median with quartiles; ****$p$<0.0001, *t* test (**C**).

**Figure S5. Aged TnC-KO mice exhibit a skeletal muscle phenotype both during homeostasis and regeneration, related to Figure 6.**

(**A**) FACS plots indicating the prevalence (%) of MuSC in whole lower hindlimb muscle samples from young versus old WT mice.

(**B**) Quantification of the wet weight (mg) of uninjured TA muscles from young and old, WT and TnC-KO mice (n≥3).

(**C**) Sirius red staining on cross-sections of uninjured TA muscles of young and old WT and TnC-KO mice (scale bar= 50 µm) and quantification of fibrosis (Sirius red signal) normalized to the young WT control.

(**D**) Representative immunofluorescence images of diaphragm cross-sections from young and old WT and TnC-KO mice (laminin, green; DAPI, blue) (scale bar= 50 µm) and quantification of the cross-sectional area (CSA, $\mu m^2$) (n=3).

(**E**) Quantification of the number of regenerating myofibers (per $mm^2$) and their CSA ($\mu m^2$) from young and old WT and TnC-KO mice (based on eMyHC staining in **Figure 6**).

Data are represented as mean ± SEM; *$p$<0.05, **$p$<0.005, one-way ANOVA (**B**, **C**, **D**, **E**).

Figure S1

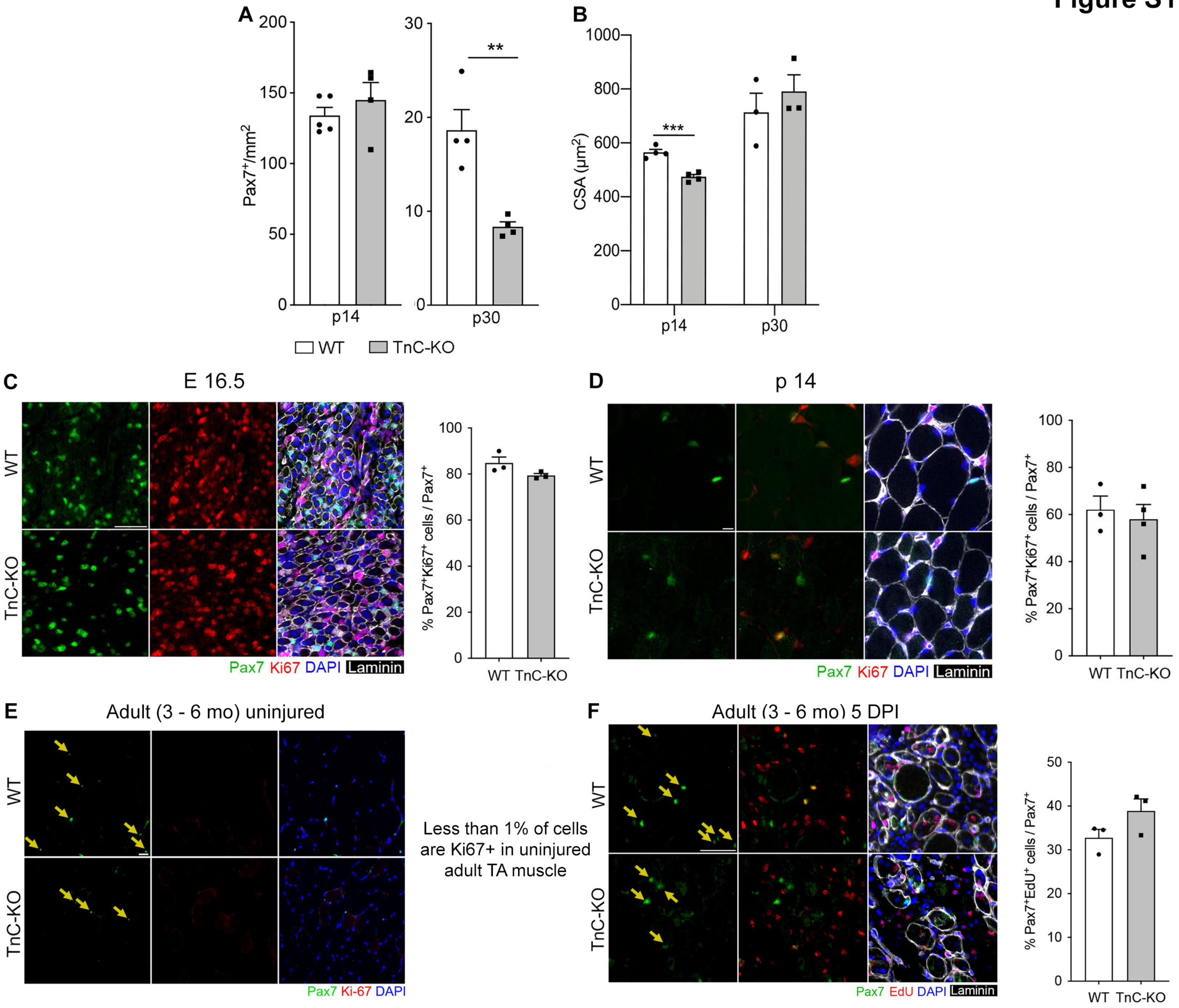

A
Pax7+/mm2
200
150
100
50
0
p14
30
20
10
0
**
p30
B
CSA (μm2)
1000
800
600
400
200
0
***
p14
p30
WT
TnC-KO
C
E 16.5
WT
TnC-KO
Pax7 Ki67 DAPI Laminin
% Pax7+Ki67+ cells / Pax7+
100
80
60
40
20
0
WT TnC-KO
D
p 14
WT
TnC-KO
Pax7 Ki67 DAPI Laminin
% Pax7+Ki67+ cells / Pax7+
100
80
60
40
20
0
WT TnC-KO
E
Adult (3 - 6 mo) uninjured
WT
TnC-KO
Pax7 Ki-67 DAPI
Less than 1% of cells are Ki67+ in uninjured adult TA muscle
F
Adult (3 - 6 mo) 5 DPI
WT
TnC-KO
Pax7 EdU DAPI Laminin
% Pax7+EdU+ cells / Pax7+
50
40
30
20
10
0
WT TnC-KO

Figure S2

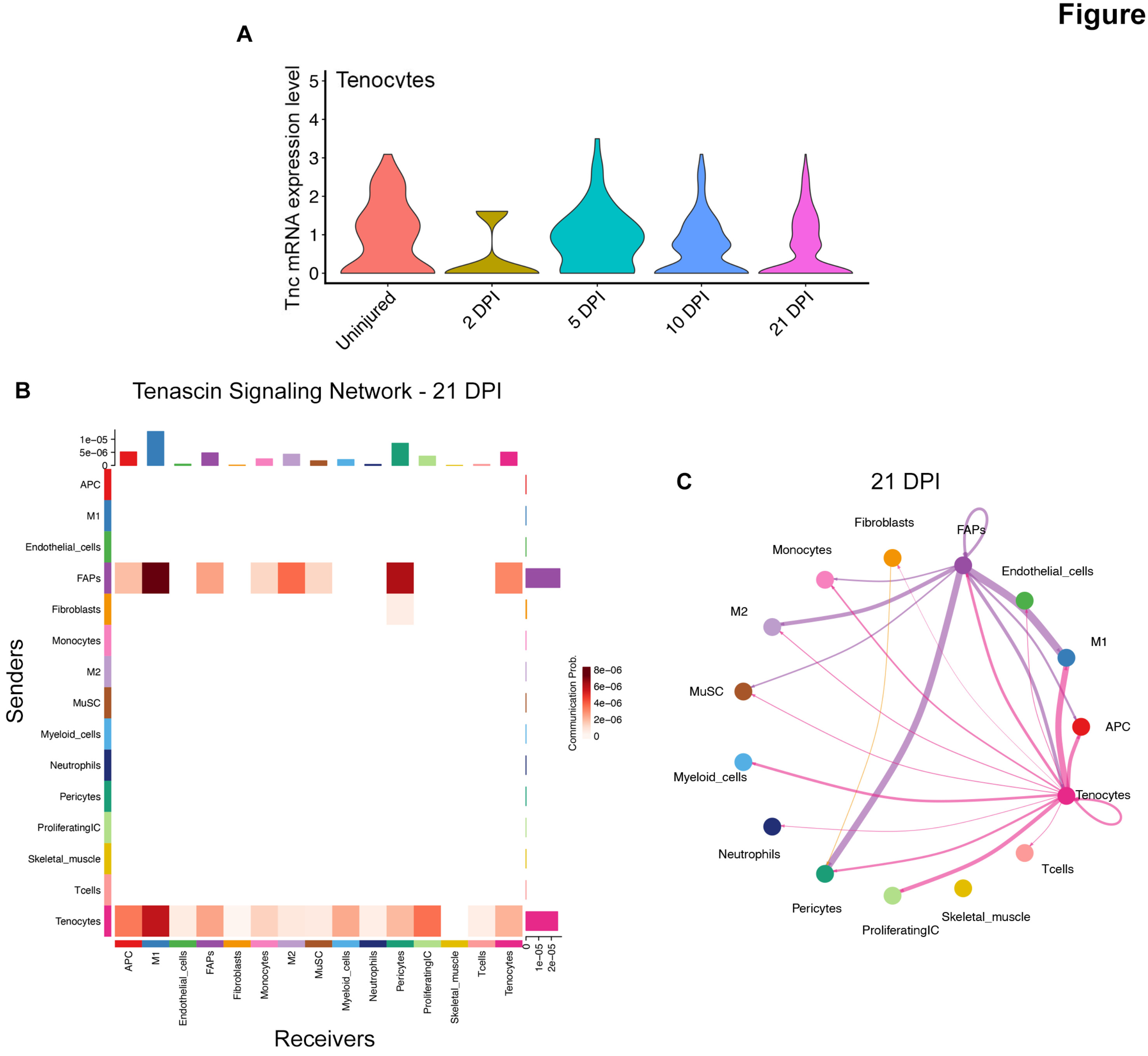
A
Tenocytes
Tnc mRNA expression level
0
1
2
3
4
5
Uninjured
2 DPI
5 DPI
10 DPI
21 DPI
B
Tenascin Signaling Network - 21 DPI
1e−05
5e−06
0
APC
M1
Endothelial_cells
FAPs
Fibroblasts
Monocytes
M2
MuSC
Myeloid_cells
Neutrophils
Pericytes
ProliferatingIC
Skeletal_muscle
Tcells
Tenocytes
Senders
Receivers
1e−05
2e−05
Communication Prob.
8e−06
6e−06
4e−06
2e−06
0
C
21 DPI
Fibroblasts
FAPs
Monocytes
Endothelial_cells
M2
M1
MuSC
APC
Myeloid_cells
Tenocytes
Neutrophils
Tcells
Pericytes
Skeletal_muscle
ProliferatingIC

**Figure S3**

**A**

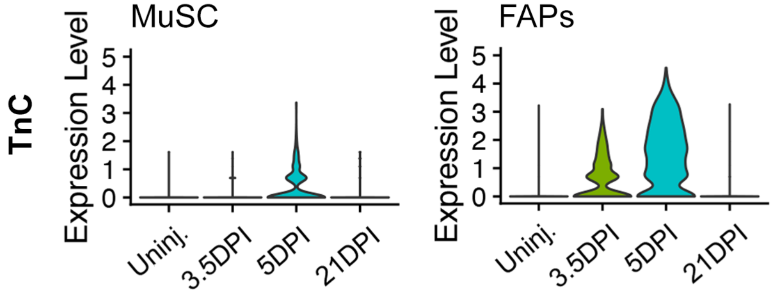


**B**

| | Uninjured | 3.5 DPI | 5 DPI | 21 DPI |
|---|---|---|---|---|
| MuSC (AnxA2+) | 104 | 235 | 304 | 61 |
| MuSC (AnxA2-) | 38 | 0 | 2 | 12 |
| FAPs (AnxA2+) | 3031 | 757 | 1103 | 3360 |
| FAPs (AnxA2-) | 50 | 10 | 19 | 154 |

**C**

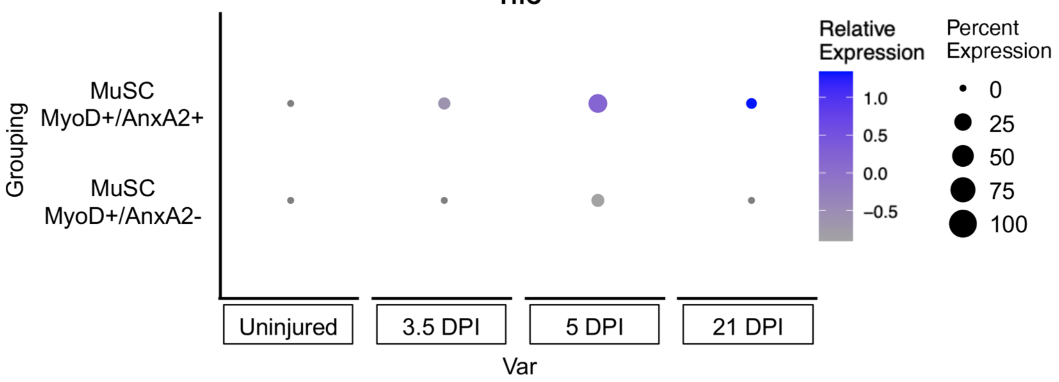


| | Uninjured | 3.5 DPI | 5 DPI | 21 DPI |
|---|---|---|---|---|
| MuSC (MyoD+/ AnxA2+) | 69 | 157 | 181 | 60 |
| MuSC (MyoD+/ AnxA2-) | 24 | 1 | 48 | 10 |

**D**

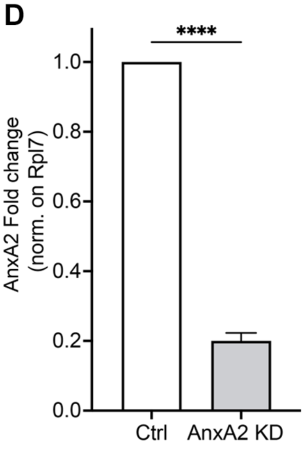


**E**

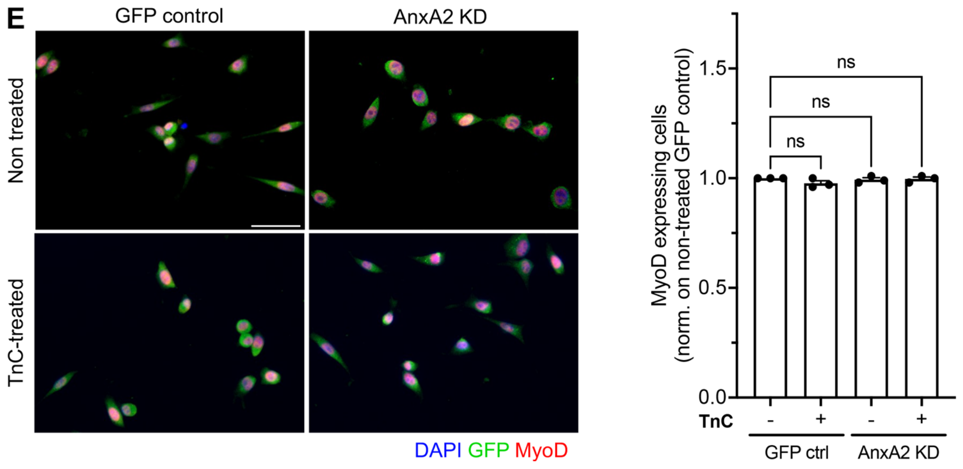

### Figure S4

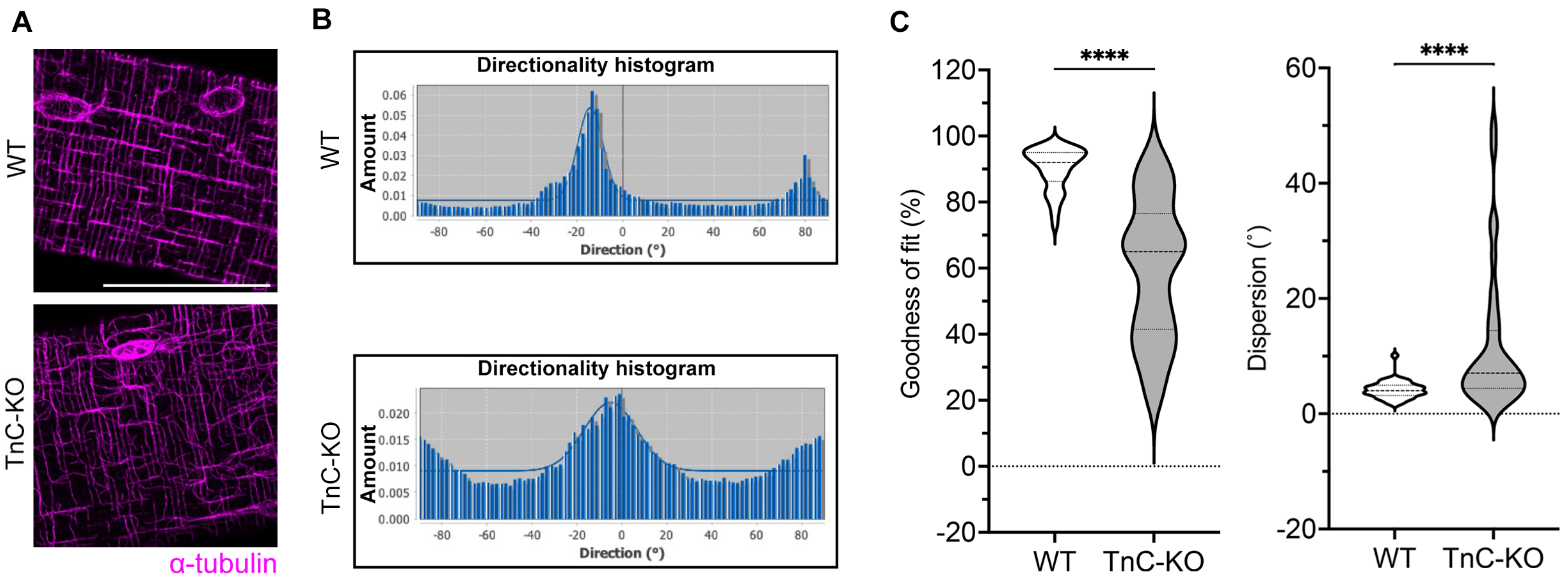

A
WT
TnC-KO
α-tubulin
B
WT
Directionality histogram
Amount
Direction (°)
TnC-KO
Directionality histogram
Amount
Direction (°)
C
****
Goodness of fit (%)
WT
TnC-KO
****
Dispersion (°)
WT
TnC-KO

### Figure S5

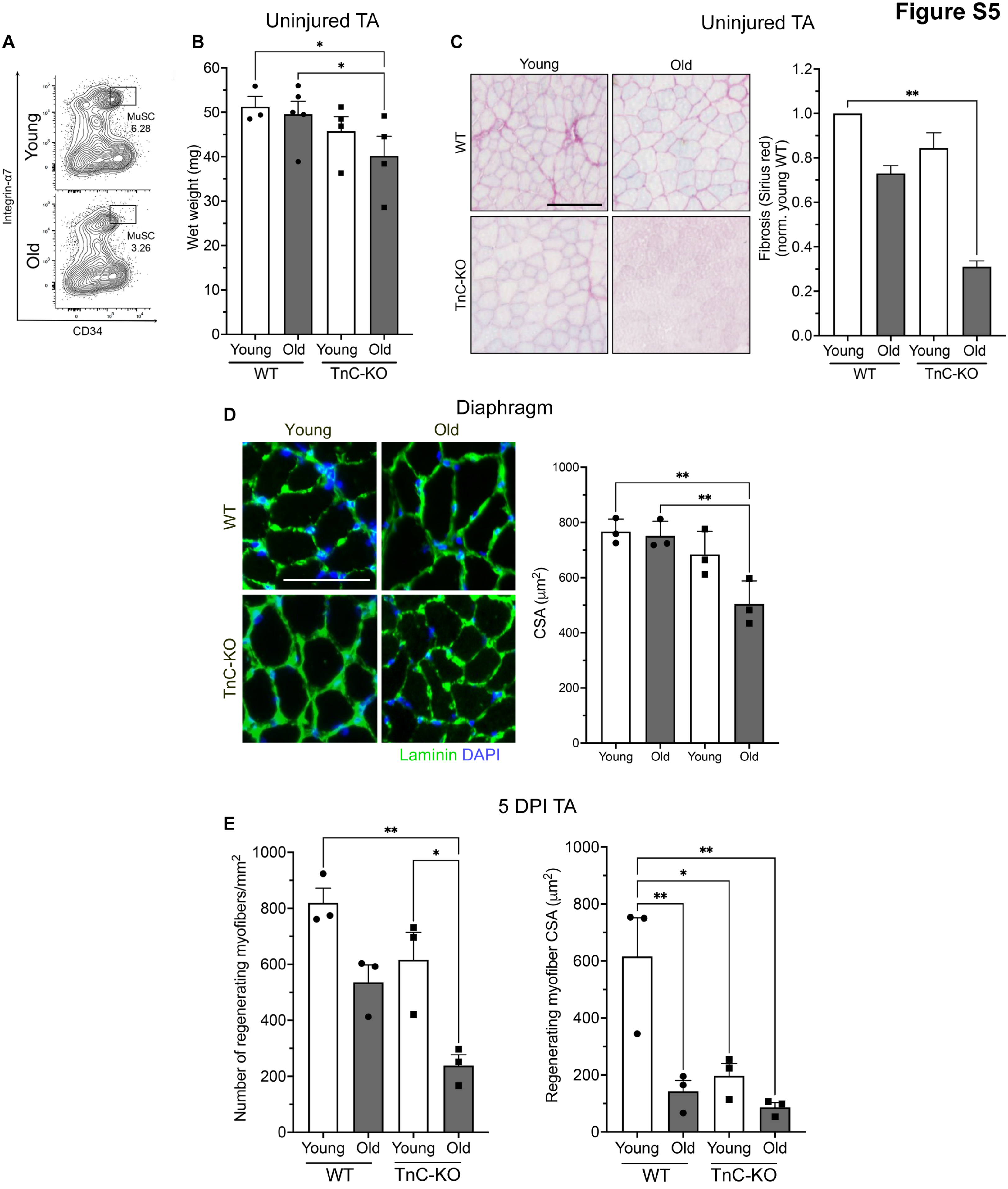
A
Young
Old
Integrin-α7
CD34
MuSC 6.28
MuSC 3.26
B
Uninjured TA
Wet weight (mg)
Young Old Young Old
WT
TnC-KO
C
Uninjured TA
Young
Old
WT
TnC-KO
Fibrosis (Sirius red) (norm. young WT)
D
Diaphragm
Young
Old
WT
TnC-KO
Laminin DAPI
CSA (μm2)
Young Old Young Old
E
5 DPI TA
Number of regenerating myofibers/mm2
Regenerating myofiber CSA (μm2)
Young Old Young Old
WT
TnC-KO
